# ACLY promotes NK cell effector function by regulating glycolysis and histone acetylation

**DOI:** 10.1101/2025.03.05.639198

**Authors:** Hyogon Sohn, Ana Kolicheski, Jennifer Laurent, Jennifer Tran, Yongjun Wang, Kelly Gan, Joshua M. Tobin, Nermina Saucier, Todd A. Fehniger, Jacqueline E. Payton, Megan A. Cooper

## Abstract

Natural Killer (NK) cells are innate immune lymphocytes important for host viral and tumor immunity. We investigated the requirement for ATP citrate lyase (ACLY) in NK cell function using an inducible genetic mouse model. ACLY regulates the citrate-malate shuttle, generating cytosolic acetyl-coenzyme A that is primarily used for acetylation or lipid synthesis. ACLY-deficient NK cells upon IL-15 activation exhibited significant defects in glycolysis, proliferation, cytokine production, and cytotoxicity, without decreased intracellular lipids. Notably, ACLY-deficiency specifically resulted in reduced NK cell responses to activating receptors associated with the adapter proteins DAP10 or DAP12. This is due to decreased DAP12 and increased DAP10 transcript and protein, coupled with epigenetic profiling that demonstrated altered histone acetylation of these genes in ACLY KO. Supplementation of ACLY-deficient NK cells with acetate was sufficient to overcome most functional defects, including restoring DAP10/12 expression and activating receptor function, emphasizing the importance of ACLY-generated cytosolic acetyl-coenzyme A for NK effector functions.

## INTRODUCTION

Natural Killer (NK) cells are innate lymphocytes that quickly respond to infections and tumor cells by producing pro-inflammatory cytokines and/or releasing cytotoxic molecules to kill target cells. NK cells are activated through germline-encoded activating and cytokine receptors that mediate their rapid response.^1^ Activating receptors recognize pathogen-encoded ligands, stress-induced molecules, the absence of a major histocompatibility complex on target cells, and the Fc region of antibodies. They transmit signals via adaptor proteins bearing immunoreceptor tyrosine-based activation motifs (ITAMs) or ITAM-like motifs, such as DAP10, DAP12, CD3ζ and FcRγ.^2–4^ For example, DAP10 and DAP12 associate with activating receptors Ly49H and NKG2D, which recognize murine cytomegalovirus (MCMV) and tumor antigens, respectively.^4–6^ Constitutively expressed cytokine receptors allow NK cells to quickly respond to inflammatory signals.^7^ Interleukin (IL)-15 in particular plays a critical role in the development and maintenance of NK cells as well as their functional responses.

Regulation of metabolic pathways and fuels is important for NK cell differentiation and function.^8–16^ NK cells exhibit distinct metabolic requirements for functional responses depending on the stimulus and the response.^14^ For example, we previously demonstrated that brief cytokine stimulation does not alter NK cell glycolysis or oxidative phosphorylation (OXPHOS), and NK cell production of IFN-γ, one of the main effector cytokines produced by NK cells, is not altered by metabolic inhibition. In contrast, stimulation of NK cells through their activating receptors is highly dependent on metabolism, and associated with post-transcriptional regulation of IFN-γ production. Culturing NK cells with high-doses of IL-15, or priming, requires glycolysis and renders NK cells resilient to metabolic perturbation.^13,15^ Priming NK cells *in vivo* also increased glycolytic capacity, induced resistant to metabolic inhibition, and exhibited efficient control of MCMV infection.^17^

Other than conventional glycolysis or tricarboxylic acid (TCA) cycle, immune cells can generate energy and intermediates through an additional catabolic process known as anaplerosis.^18^ The citrate-malate shuttle (CMS) exemplifies such a process, which begins with the translocation of mitochondrial citrate to the cytosol. In the cytosol, ATP citrate lyase (ACLY) converts the citrate to acetyl-coenzyme A (CoA) and oxaloacetate; oxaloacetate can be recycled in the mitochondria as malate and continue to fuel OXPHOS (Supplementary Figure 1A).^19^ Notably, it is known in cancer cells that ACLY-generated cytosolic acetyl-CoA is predominantly involved in histone acetylation, a canonical chromatin modification that positively regulates gene expression.^20^ Cytosolic acetyl-CoA can also be used for cytosolic protein acetylation and lipid synthesis.^18,19^

Interestingly, some studies have indicated a correlation between cytosolic acetyl-CoA levels and function of immune cells. For instance, CD4^+^ T helper 1 cells with elevated acetyl-CoA levels have been shown to promote histone acetylation at the *Ifng* promoter and CNS22 enhancer region.^21^ Moreover, cytosolic acetyl-CoA mediates CD8^+^ T cell differentiation and functional responses to pathogens *in vivo*.^22,23^ In NK cells, it was previously shown that cytokine stimulation with IL-2/12 activates the transcription factor SREBP, which rewires cell metabolism to upregulate CMS and ACLY activity.^8^ However, the metabolic-epigenetic axis in NK cells is not fully understood.

Here, we generated a genetic mouse model with conditional deletion of *Acly* in NK cells and report that during IL-15 priming ACLY is required for upregulating glycolysis, NK cell proliferation, cytotoxicity and responses to activating receptors that partner with DAP10 and DAP12.

## RESULTS

### Inducible deletion of *Acly* in NK cells does not alter NK cell phenotype at baseline

To investigate the connection between metabolism and the epigenome in NK cells, we generated a tamoxifen-inducible model of NK-specific *Acly* deficiency (referred to as ‘ACLY KO’ here). NK cells with *Acly*-deficiency were also marked by YFP expression by crossing to a reporter mouse (*Ncr1*^iCre-ERT2^*Rosa*^YFP/YFP^*Acly*^fl/fl^, see methods section), and cells were gated on YFP to confirm Cre activity. Following tamoxifen administration in the chow, YFP^+^ NK cells lacked expression of *Acly* mRNA and protein (Supplementary Figure 1B-D and Supplementary Figure 7A-B), whereas the small fraction of remaining YFP^-^ NK cells still produced ACLY protein. Next, we compared total cell numbers between WT (*Ncr1^iCre-ERT2^Rosa^YFP/YFP^* controls) and ACLY KO mice following tamoxifen treatment. The absolute number of YFP^+^ NK cells in the spleen, liver, blood and bone marrow (BM) were not affected by acute ACLY deletion (Supplementary Figure 1E).

NK cell phenotype in multiple organs (spleen, liver, blood, and BM) was examined to assess the impact of induced *Acly* deletion on NK cells. *Acly* deletion did not impact the expression of maturation markers CD27 and CD11b in NK cells, as the percentage across the four-populations (CD11b^-^ CD27^-^, CD11b^-^ CD27^+^, CD11b^+^ CD27^+^, CD11b^-^ CD27^+^) remained similar (Supplementary Figure 1F). Expression of activating and inhibitory NK receptors was measured by flow cytometry, and YFP^+^ NK cells from ACLY KO mice and WT mice had similar expression levels for all examined receptors (Ly49A, Ly49D, Ly49H, CD122, NKG2D, NK1.1, CD132, NKp46) (Supplementary Figure 1G-H). Thus, with acute *Acly* deletion there was no apparent impact on the abundance of NK cells or the expression of tested receptors.

In CD4 T cells, ACLY supports proliferation by providing acetyl-CoA for fatty acid synthesis and histone acetylation.^24^ Thus, we hypothesized that acute deletion of ACLY might alter proliferation of NK cells. To test this, YFP^+^ cells from congenic WT (CD45.1) and ACLY KO (CD45.2) mice were mixed at similar ratios and adoptively transferred into *Rag2*-γc KO mice, which lack T, B, and NK cells (Supplementary Figure 2A). The percentages of WT (CD45.1) and KO (CD45.2) remained similar, suggesting that the absence of ACLY does not alter NK cell homeostatic proliferation (Supplementary Figure 2B-C).

If the CMS is active, the absence of ACLY would be expected to lead to accumulation of its substrate, citrate, which inhibits phosphofructokinase-1, a key enzyme of the glycolytic pathway. In addition, NAD^+^ is a product of the ACLY-driven CMS, and promotes glycolysis. Thus, deletion of ACLY would also be expected to inhibit glycolysis if the CMS is active (Supplementary Figure 1A).^18,25,26^ Therefore, we examined whether freshly isolated ACLY KO NK cells have impaired glycolysis compared to WT by extracellular flux analysis (Seahorse^TM^ assay). In freshly isolated cells, the glycolytic rate and capacity measured by extracellular acidification rate (ECAR) was comparable between WT and KO cells, suggesting that KO cells maintain a normal glycolytic capacity under homeostatic conditions (Supplementary Figure 2D). This is consistent with prior data suggesting that the CMS is only upregulated with activation of NK cells.^8^

It was previously reported that inhibition of ACLY impaired NK cell IFN-γ production.^8^ However, in contrast to inhibitor data, freshly isolated naïve ACLY KO NK cells have normal IFN-γ responses with cytokine or receptor stimulation (Figure 1A and Supplementary Figure 2E). Furthermore, treatment with the ACLY inhibitor used in this prior report and other immunometabolism studies (ACLYi, BMS-303141), significantly impaired IL-2 + IL-12 induced IFN-γ expression by both WT and ACLY KO cells (Figure 1B). Interestingly, when NK cells were pre-cultured with low-dose IL-15 for 6 days, similar to ref^8^, there was significant reduction in IFN-γ production in ACLY KO but not WT cells. Together these studies indicate off-target effects of this ACLY inhibitor.

**Figure 1.**
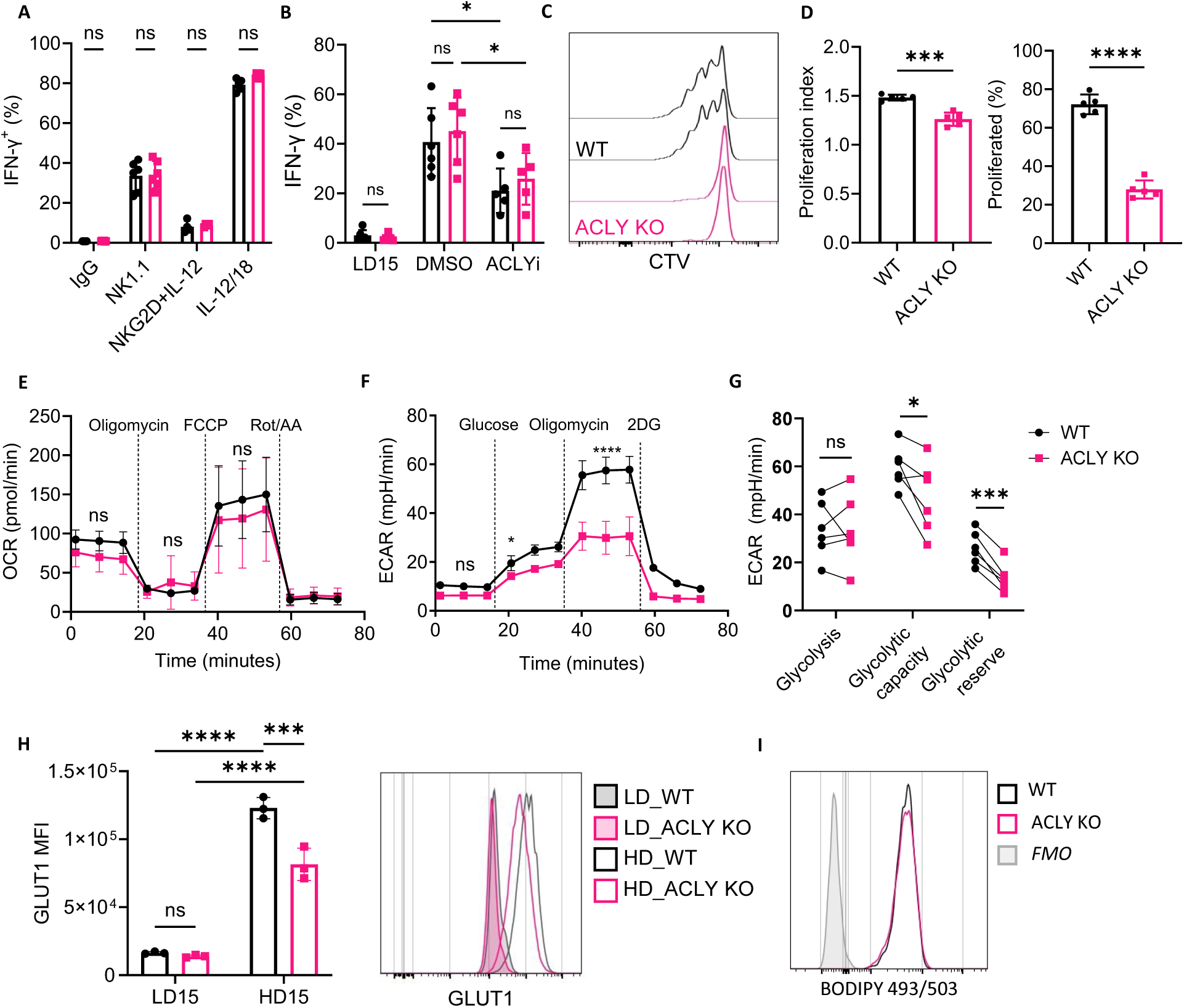
Primed *Acly*-deficient NK cells have proliferation defects with diminished glycolytic capacity. Splenic NK cells from *Ncr1*^iCre-ERT2^*Rosa*^YFP/YFP^ (WT) and *Ncr1*^iCre-ERT2^*Rosa*^YFP/YFP^*Acly*^fl/fl^ (ACLY KO) were analyzed *in vitro*, using YFP expression to identify cells with Cre activity. Spleens were analyzed 5 days after withdrawing tamoxifen diet. (**A and B**) IFN-γ expressing YFP^+^ NK cells were analyzed by intracellular flow cytometry. Freshly isolated NK cells stimulated with anti-NK1.1, anti-NKG2D + IL-12 (10ng/ml), or IL-12 (10ng/ml) + IL-18 (50 ng/ml) for 6 hours (A), or with IL-2 (20 ng/ml) and IL-12 (10 ng/ml) in the presence of DMSO (carrier) or the ACLY inhibitor BMS-303141 (50μM) for 18 hours (B). Results represent 2 independent experiments with n=3-6 biological replicates, two-way ANOVA. (**C through G**) WT or ACLY KO NK were primed with high-dose (HD) IL-15 (100ng/mL) *in vitro* for 72 hours. (**C and D**) NK cells were labeled with cell trace violet (CTV) prior to priming to evaluate proliferation, with representative CTV dilution of YFP^+^ NK cells (C) and proliferation index and percentage proliferated cells shown (D). Results represent 3 independent experiments with n=5 biological replicates, unpaired t-test. (**E through G**) Extracellular flux analysis of sorted and IL-15 primed YFP^+^ WT or ACLY KO NK cells with drugs shown to test oxidative stress by oxygen consumption rate (OCR) (E) and glycolytic capacity with extracellular acidification rate (ECAR) shown (F-G). Representative data of 3 (E) or 6 (F) independent experiments, two-way ANOVA (G) Summary of glycolysis, glycolytic capacity, and glycolytic reserve from 6 independent experiments. Paired t-test. (**H**) MFI (left) and representative flow histogram (right) of glucose transporter (GLUT1) by flow cytometry. Results represent 2 independent experiments with n=3 biological replicates, two-way ANOVA. (**I**) Representative flow histogram of BODIPY 493/503 staining in primed YFP^+^ NK cells. Shaded gray indicates fluorescence minus one (FMO) control. Data are shown as mean ± SD, *p≤0.05, **p≤0.01, ***p≤0.001, ****p≤0.0001

Together, these experiments demonstrate that acute deletion of *Acly* in NK cells does not significantly impact receptor expression, proliferation, glycolytic capacity, or IFN-γ response to cytokines or activating receptors. This genetic model also demonstrates off-target effects of a commonly used ACLY inhibitor.

### *Acly*-deficiency impairs proliferation and glycolysis of NK cells with IL-15 priming

IL-15 is a key cytokine for NK cells, supporting proliferation, survival, and cytokine production *in vivo*.^13,14,27^ Notably, varying levels of IL-15 can have differing effects on NK cells: low doses of IL-15 (10 ng/mL) promote NK cell survival, whereas higher concentrations of IL-15 (100 ng/mL; HD15) drive NK cell activation, proliferation, cytotoxicity and reprograms metabolism by upregulating both glycolysis and OXPHOS.^11,14,15,28,29^ It was previously shown that cytokine-activated NK cells upregulate the transcription of *Srebp1* and *Srebp2* (encoding transcription factor SREBP) leading to upregulation of *Acly* expression.^8,30,31^ We hypothesized that at steady-state metabolic reserve, ACLY may not be important for NK cell function, but with priming and activation of this pathway may be critical.

Indeed, IL-15 priming of ACLY KO cells led to a significant proliferation defect (Figure 1C-D). With rapid proliferation, NK cells undergo a metabolic switch from OXPHOS to glycolysis.^15^ While primed ACLY KO NK cells exhibited normal mitochondrial respiration (Figure 1E), they had significantly reduced glycolytic capacity and spare glycolytic reserve (Figure 1F-G) in extracellular flux assays. Based on this defect, we tested expression of the primary glucose transporter used by NK cells, GLUT1, by flow cytometry. Both WT and ACLY KO NK cells upregulated GLUT1, however primed WT NK cells expressed significantly higher levels of GLUT1 than ACLY KO cells (Figure 1H).

However, fatty acids can support NK cell proliferation *in vitro* and *in vivo*, and ACLY-derived cytosolic acetyl-CoA is a substrate for lipid synthesis (Supplementary Figure 1A).^32,33^ To test whether the absence of ACLY caused a defect in lipid synthesis, we compared the level of intracellular neutral lipids between primed WT and ACLY KO cells using BODIPY staining. This demonstrated no difference (Figure 1I), suggesting that deficiency of *Acly* does not alter neutral lipid contents in primed NK cells.

Together, these results demonstrate that ACLY is not required for NK cell function at steady-state, but is required for the increased metabolic demands of activation and priming, including upregulation of glycolysis and cellular proliferation.^34^

### Primed ACLY KO NK cells show specific defects in response to NKG2D and Ly49H stimulation

Primed NK cells have robust responses to stimulation of NK activating receptors including NK1.1, Ly49H, and NKG2D in the C57BL/6 mouse.^13^ This effect of priming is independent of proliferation, as undivided cells and highly divided cells respond similarly.^15^ Moreover, primed NK cells obtain metabolic flexibility *in vivo* and *in vitro* and their IFN-γ production is no longer dependent on glycolysis or OXPHOS.^13,15,17^

IL-15 primed (HD15) WT and ACLY KO NK cells were tested for their response to activating receptor stimulation (anti-NK1.1, anti-NKG2D and anti-Ly49H) compared to unstimulated cells cultured for the same duration period with LD15 (low-dose IL0-15) to maintain survival. Primed WT and ACLY KO NK cells expressed similarly high levels of IFN-γ following NK1.1 stimulation (Figure 2A). However, primed ACLY KO cells had a significant defect in IFN-γ production when stimulated via NKG2D or Ly49H, with levels similar to LD15 treated cells (Figure 2B-C). Priming did not alter the already high NK cell IFN-γ responses to PMA and Calcimycin (PMA/Cal), and WT and ACLY KO NK cells had similar responses with LD15 or HD15 (Figure 2D).

**Figure 2.**
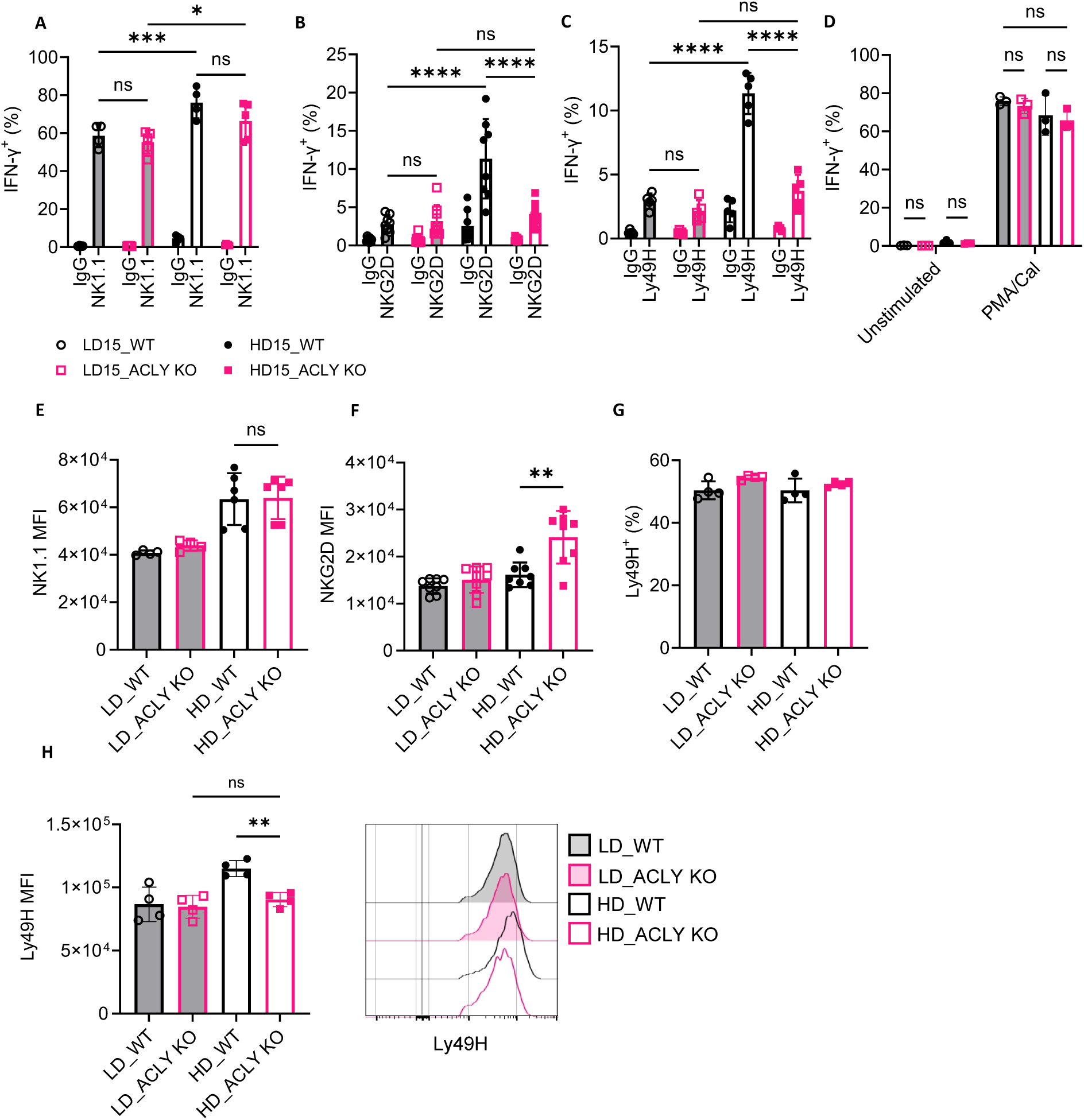
Primed *Acly*-deficient NK cells have impaired response to activating receptors. (**A through C**) NK cells were cultured in LD15 (10ng/ml IL-15) or HD15 (100ng/ml IL-15) *in vitro* for 72 hours. IFN-γ expressing YFP^+^ NK cells were measured after stimulated with IgG, NK1.1 (A), NKG2D (B), or Ly49H (C) for 6 hours by intracellular staining. Results represent 3-5 independent experiments with n=4-8 biological replicates, two-way ANOVA. (**D**) NK cells were cultured in LD15 or HD15 *in vitro* for 72 hours and stimulated with PMA/Calcimycin for 6 hours. Percentage of IFN-γ expressing YFP^+^ cells were analyzed by intracellular flow cytometry. Results represent 2 independent experiments with n=3 biological replicates, two-way ANOVA. (**E and F**) Mean Fluorescence Intensity (MFI) of NK1.1 (E) and NKG2D (F) on non-primed (LD) or primed (HD) YFP^+^ NK cells. Results represent 3-5 independent experiments with n=6-8 biological replicates, unpaired t-test. (**G and H**) Percentage of Ly49H^+^ population (G) and MFI with representative histogram (H) of Ly49H expression among YFP^+^ NK cells. Results represent 4 independent experiments, unpaired t-test. Data are shown as mean ± SD, *p≤0.05, **p≤0.01, ***p≤0.001, ****p≤0.0001

Given the observed defects, cell-surface expression of the tested activating receptors was measured. All NK cells expressed NK1.1, and the mean fluorescence intensity (MFI) of this receptor increased after priming in both WT and ACLY KO NK cells (Figure 2E). Interestingly, primed ACLY KO NK cells had higher cell surface density of NKG2D compared to WT NK cells (Figure 2F). While the percentage of NK cells expressing Ly49H was similar between WT and ACLY KO NK cells, there was a small but significant decrease in Ly49H MFI in ACLY KO NK cells, which could cause the defects in cytokine production (Figure 2G-H).

Together, these findings indicate that ACLY deficient primed NK cells have the capacity for high IFN-γ protein production, but that their response to certain activating receptors, specifically Ly49H and NKG2D, is impaired.

### ACLY is critical for cytotoxicity of primed NK cells

We and others have previously demonstrated that upregulation of glycolysis is essential for NK cell killing including in the setting of viral infection and tumors *in vivo.*^17,35^ Priming with IL-15 induces a shift to glycolysis in NK cells and translation of key cytotoxic proteins, granzyme B (Gzmb) and perforin (Prf1).^9,29^ Based on the observed impaired glycolysis with *Acly* deficiency, we hypothesized that NK cell killing would be affected. Both primed WT and ACLY KO cells had increased intracellular Prf1 and Gzmb proteins compared to cells cultured with LD15; however, ACLY KO cells had significantly lower levels of Prf1 compared to primed WT NK cells (Figures 3A-B). NK cell recognition of target cells and degranulation was measured by upregulation of CD107a in primed WT and ACLY KO cells following culture with the NK-resistant target RMA, or RMA cells overexpressing the NKG2D ligand Rae1 (RMA-Rae1) or Ly49H ligand m157 (RMA-m157). Both WT and KO cells had minimal degranulation with RMA, with higher levels in response to RMA-Rae1 and RMA-m157 cells (Figure 3C-D). ACLY KO cells exhibited similar degranulation to WT in response to RMA-Rae1 but showed reduced degranulation in response to RMA-m157, which could be due to the lower cell-surface density of Ly49H or to impaired Ly49H downstream signaling.

**Figure 3.**
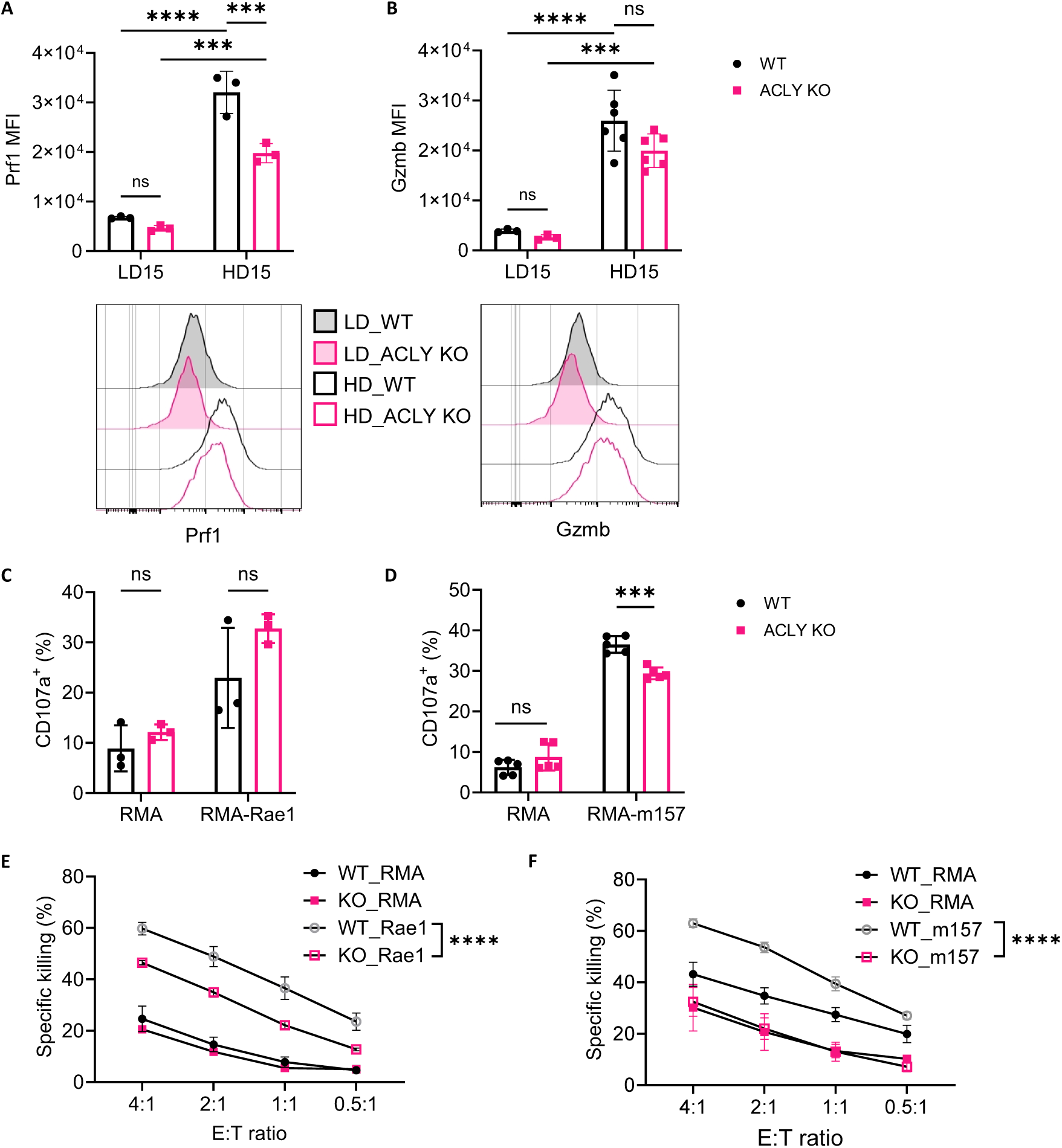
Impaired NK cell cytotoxic function with *Acly* deficiency. NK cells were primed with HD15 *in vitro* for 72 hours. (**A and B**) MFI and representative histogram of perforin (A) and granzyme B (B) in YFP^+^ NK cells cultured in LD15 or HD15. Result represents 2-3 independent experiments with n=3-6 biological replicates, two-way ANOVA. (**C and D**) Sorted YFP^+^ cells were mixed in a 1:1 ratio with either RMA or RMA-Rae1 (C) and enriched NK cells were mixed in a 1:1 ratio with RMA or RMA-m157 cells (D), followed by a 6-hour incubation. Percentage of CD107a on YFP^+^ NK cells was measured by flow cytometry. Results represent 1 (C) and 3 (D) independent experiments with n=3 biological replicates each, two-way ANOVA. (**E and F**) YFP^+^ NK cells (effectors) were sorted 3 days after tamoxifen treatment, primed with IL-15, and co-cultured with CTV^+^ RMA-Rae1 (D) or RMA-m157 (E) targets in effector to target (E:T) ratios shown. Percent specific killing was analyzed by flow after 4 hours. Results represent 1-2 independent experiments with n=3 biological replicates each, mean ± SEM, two-way ANOVA. Data are shown as mean ± SD, *p≤0.05, **p≤0.01, ***p≤0.001, ****p≤0.0001

Degranulation of cytotoxic molecules is required for target cell killing, therefore we measured NK cell killing using a flow-based *ex vivo* killing assay with primed NK cells and target cells mixed in various ratios. As expected, WT cells had enhanced killing of RMA-Rae1 and RMA-m157 cells compared to RMA However, ACLY KO NK cells had significantly lower killing compared to WT (Figure 3E-F). Although ACLY KO NK cells exhibited similar degranulation and higher receptor expression levels, reduced level of Prf1 significantly affected the cytotoxicity of NK cells.

These results suggest that primed NK cells lacking ACLY have reduced levels of cytotoxic molecules, thereby reducing their cytotoxicity capabilities against target cells.

### ACLY KO NK cells have an altered transcriptional profile in response to IL-15 priming and NKG2D stimulation

To explore how genetic deletion of *Acly* impairs primed NK cell function, we performed whole transcriptome analysis (RNA-seq) on YFP^+^ fresh, HD15 (primed), and NKG2D-stimulated primed WT and ACLY KO NK cells (Supplementary Figure 3A). There were minimal transcriptional differences between fresh WT and ACLY KO, with the most significantly elevated transcript in WT NK cells being *Acly*, consistent with the genetic model (Supplementary Figure 3B). There were more significant differences between primed WT and KO NK cells without and with NKG2D stimulation (Figure 4A and B respectively). Consistent with protein data, ACLY KO primed NK cells expressed lower level of *Ifng* transcript than WT after NKG2D stimulation (Figure 4B and C). Transcripts for additional cytokines and chemokines downregulated in ACLY KO NK cells, include *Ccl3*, *Cx3cl1*, *Tnf* following NKG2D stimulation. ACLY KO NK cells also had impaired upregulation of genes encoding cytotoxic molecules including transcripts for granzymes and perforin, with IL-15 priming and after NKG2D stimulation (Figure 4A-C).

**Figure 4.**
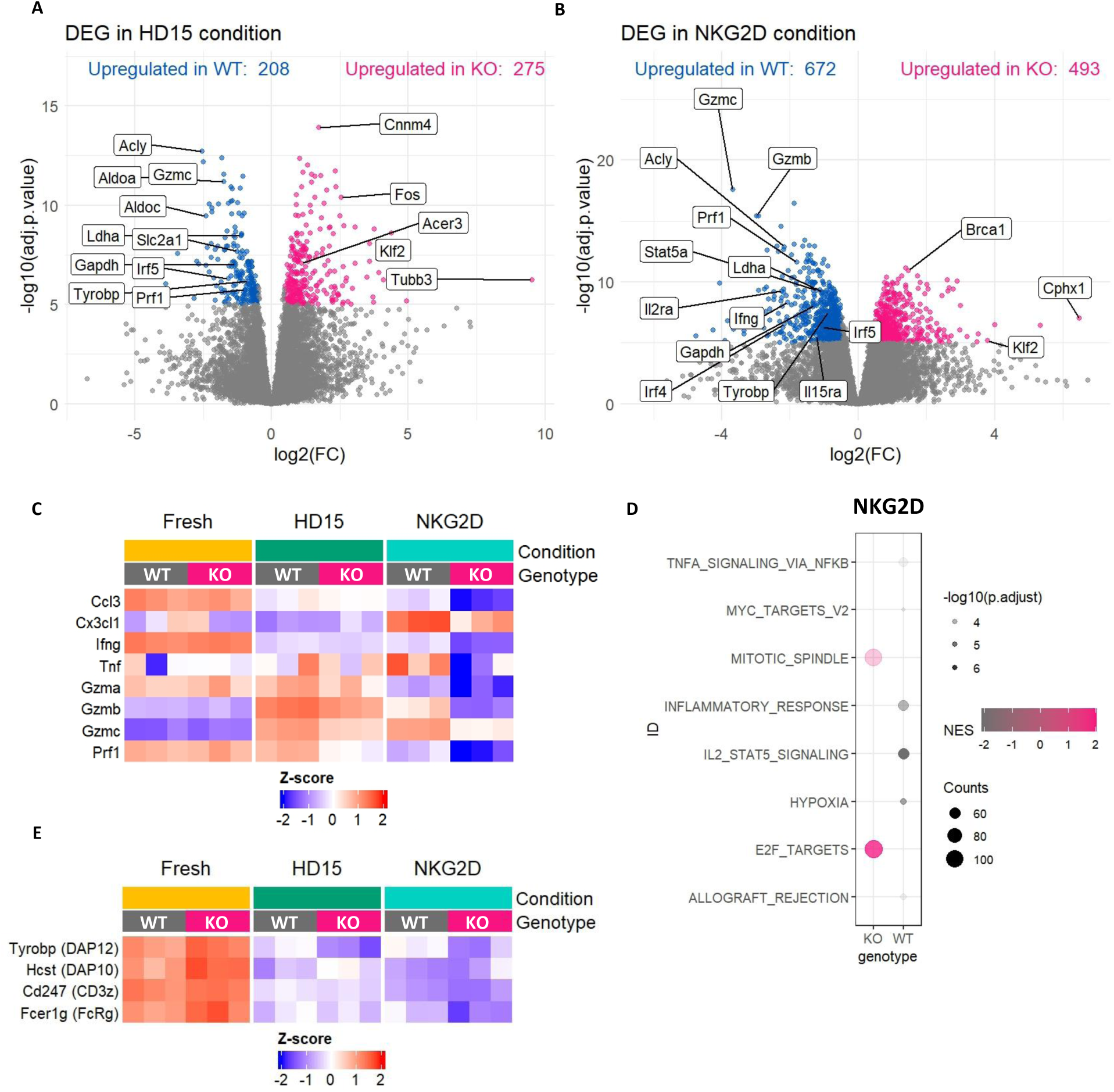
Transcriptional profiling of *Acly*-deficient NK cells demonstrates differential gene expression. YFP^+^ NK cells from the spleen of 2 mice were pooled for each sample. Results represent 3 biological replicates per condition and genotype (see also Supplementary Figure 3). (**A and B**) Volcano plots showing differentially expressed genes (DEG) between WT and ACLY KO NK cells after HD15 priming (A) or priming followed by NKG2D stimulation (B). (**C**) Heatmap of selected genes related to cytokines and cytotoxic molecules that were differentially expressed between WT and ACLY KO in HD15 or/and priming + NKG2D conditions. (**D**) Bubble plot of gene set enrichment analysis (GSEA) of Hallmark pathways upregulated (magenta) and downregulated (gray) in primed ACLY KO as compared to WT NK cells after NKG2D stimulation. (**E**) Heatmap showing transcriptional changes of selected genes (with corresponding protein names) in Fresh, HD15, and NKG2D conditions.

Pathway analysis (gene set enrichment assay, GSEA) demonstrated that ACLY KO cells fail to upregulate genes related to glycolysis and hypoxia, such as *Gapdh*, *Hk2*, *Ldha*, and *Slc2a1* (Figure 4A and Supplementary Figure 3C) during priming, consistent with metabolic profiling results (Figure 1F). This transcriptional data suggests that ACLY KO cells failed to prepare the metabolic machineries for NK cell priming, which are essential for cell proliferation.^11,17^ Following NKG2D stimulation, primed ACLY KO cells consistently failed to upregulate hypoxia genes (Figure 4B and D). In addition, in WT but not ACLY KO NK cells, NKG2D stimulation strongly upregulated STAT5 targets, inflammatory response genes and NFκB targets (Figure 4D). To elucidate genes that were specifically upregulated by NKG2D stimulation, we conducted differential expression analysis between NKG2D and HD15 in each genotype. Genes that had log_2_ fold change (FC) > 0.5 and adjusted p-value < 1e-5 were called significant. WT and ACLY KO NK cells only shared 30.4% of genes that were upregulated by NKG2D stimulation (Supplementary Figure 3D), demonstrating altered NKG2D signaling in ACLY KO NK cells.

Primed ACLY KO cells upregulated cell-surface expression of NKG2D (Figure 2F) and we further investigated whether NKG2D expression was upregulated from transcript level. There are two isoforms of murine NKG2D, long (232 aa) and short (219 aa), which arise from alternative splicing of the *Klrk1* transcript.^6,36^ Based on RNAseq data, NKG2D-S (short) expression increased upon IL-15 priming in both WT and KO. In contrast, NKG2D-L (long) expression was downregulated with priming in WT but remained unchanged in KO cells (log_2_FC = 1.05) (Supplementary Figure 3E). As a result, KO cells exhibited increased NKG2D transcript suggesting that increased cell-surface protein expression is transcriptionally regulated in ACLY KO NK cells.

Overall, RNA-seq demonstrated that ACLY plays a critical role in upregulating the expression of genes involved in glycolysis pathways and effector molecules in primed NK cells. Upon NKG2D stimulation, failure to upregulate STAT5 targets correlate with impaired cytokine expression in ACLY KO NK cells.

### Defects in DAP10/12 regulation in ACLY KO NK cells

Having observed that ACLY KO NK cells have a selective defect in response to NKG2D and Ly49H, but not other pathways including anti-NK1.1, cytokines and PMA/Calcimycin, we focused on factors common to these activating receptor pathways. Both NKG2D and Ly49H signaling depend on adaptor proteins DAP10 or DAP12, which each can bind to the intracellular domain of NKG2D and Ly49H, and association with either adaptor is required for cell-surface expression and signaling of these receptors (Supplementary Figure 4A).^5,6^ By contrast, NK1.1 utilizes the adaptor protein FcRγ.^37^ Upon receptor ligation, DAP10 and DAP12 both transduce activating signals, but they are not interchangeable and have distinct functions in NK cells. DAP12 contains an ITAM motif that activates Syk/Zap70 and is thought to primarily be important for cytokine production by NK cells.^4,38^ In contrast, DAP10 has a YINM motif and transduces signals through Erk and PLCγ2 to trigger target killing and augment signals initiated through other receptors in CD8^+^ T cells.^4,5,39^ Distinct from humans, different isoform of mouse NKG2D can associate with DAP10 or DAP12. NKG2D-L is associated with DAP10 for surface presentation in resting state, whereas NKG2D-S can bind to both DAP10 and DAP12 and is upregulated in activated NK cells.^4,6,36,39^ Based on impaired NKG2D stimulation despite higher cell-surface expression of this receptor, we hypothesized that DAP10 and DAP12 signaling may be altered.

RNA-seq data was further analyzed for expression of 4 main adaptor proteins in NK cell; *Tyrobp* (encodes DAP12), *Hcst* (encodes DAP10), *Cd247* (encodes CD3ζ), and *Fcer1g* (encodes FcRγ) (Figure 4E). The expression of all genes was downregulated upon IL-15 priming, possibly due to negative feedback regulatory mechanisms. However, with HD15 priming, ACLY KO NK cells expressed lower levels of *Tyrobp* (DAP12) (log_2_FC = –0.87) and higher *Hcst* (DAP10) (log_2_FC = 0.43) compared to WT cells, whereas the levels of other NK adaptor protein transcripts (CD3ζ, FcRγ) were similar (log_2_FC = – 0.08 and –0.07 respectively). This also translated to the protein level, with significantly higher levels of DAP10 and lower DAP12 protein in primed ACLY KO cells by immunoblot (Figure 5A-B and Supplementary Figure 7C-E). Naïve NK cells had similar mRNA expression of *Tyrobp* and *Hcst* and the encoded proteins (DAP12 & DAP10 respectively), suggesting that varying expression of adaptor protein was the result of IL-15 priming in WT and ACLY KO cells (Figure 4E, Supplementary Figure 4B-C, and Supplementary Figure 7F-G). We hypothesize that ACLY KO NK cells compensate for low levels of DAP12 by upregulating DAP10, allowing NKG2D and Ly49H to be presented on the cell surface but at the cost of less ITAM signal transduction upon activation.

**Figure 5.**
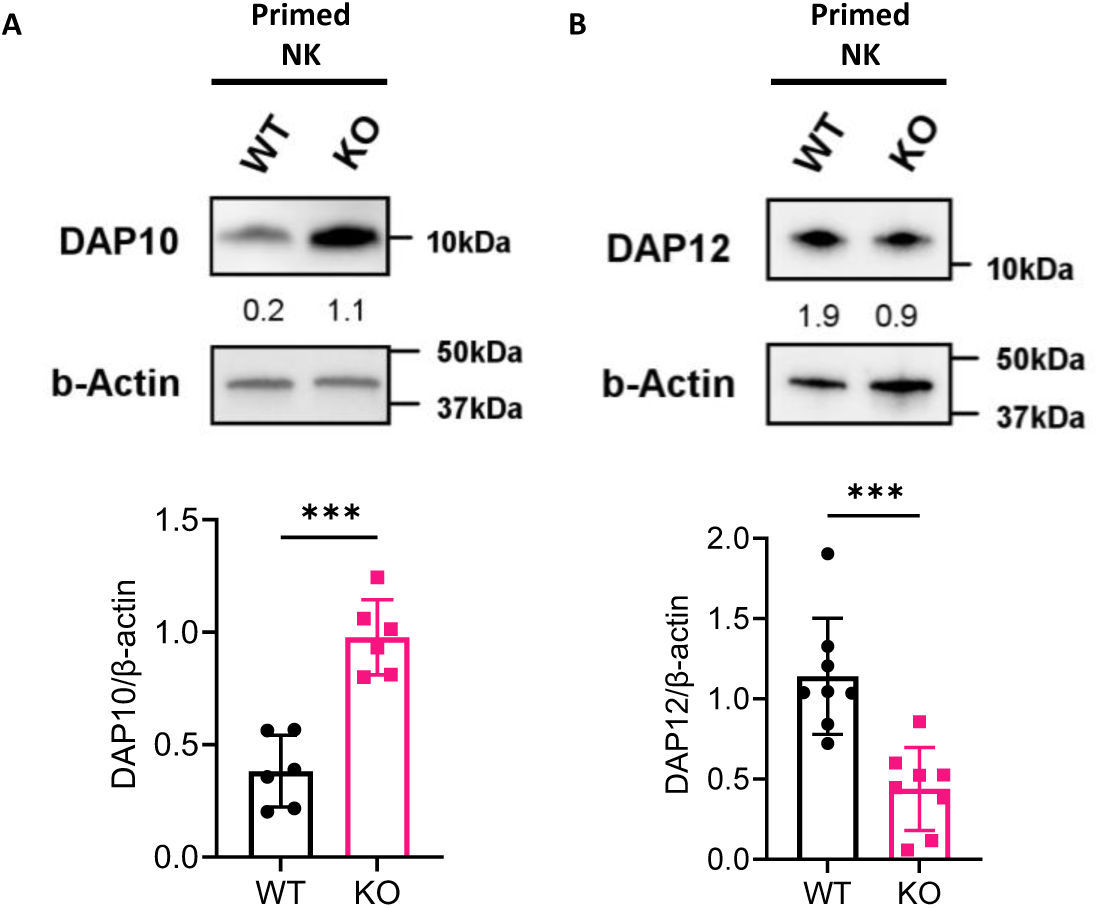
*Acly*-deficiency alters the expression of DAP10 and DAP12 protein. (**A and B**) Sorted YFP^+^ WT and ACLY KO NK cells were primed with HD15 for 72 hours. Protein expression of DAP10 (A) and DAP12 (B) were examined by immunoblotting. Protein intensity was quantified by normalizing the intensity of the protein of interest to that of β-actin. Bar graph shows the relative expression levels of proteins. Results represent with n=6-8 biological replicates, paired t-test. See Figure S7C-E for original image. Data are shown as mean ± SD, *p≤0.05, **p≤0.01, ***p≤0.001, ****p≤0.0001

### ACLY is required for genome-wide epigenomic remodeling during IL-15 priming

Cytosolic acetyl-CoA generated by ACLY can serve as the substrate for histone acetylation, which is associated with activated chromatin states and increased gene transcription (Supplementary Figure 1A).^19,20,22,40–43^ We hypothesized that higher levels of histone acetylation could be responsible for varying transcription levels of DAP10 and DAP12. To test this, the abundance of active acetylation (H3K27ac, H3K9ac), and repressive methylation (H3K27me3) chromatin marks were compared between primed WT and ACLY KO NK cells using Cleavage Under Targets and Tagmentation (CUT&Tag) sequencing.^44^ H3K27ac and H3K9ac are histone acetylation marks that define active promoters and enhancers, but H3K27ac is specifically associated with active enhancers.^45,46^ Differentially binding analysis was performed using DiffBind through DESeq2. Peaks were considered significant if they had an FDR < 0.05 and an absolute log_2_FC ≥ 0.5. As a result, ACLY KO cells exhibited 1611 peaks with reduced H3K27ac marks and 1590 peaks with increased H3K27ac marks (Figure 6A and Supplementary Figure 5A). The decreased H3K27ac peaks were near genes related to glycolysis (*Rptor*, *Ldha*), Il-15 receptor (*Il15ra*) and *Nfkb1*. In addition, 785 peaks showed reduced H3K9ac marks, while 352 peaks exhibited increased H3K9ac marks in ACLY KO cells (Figure 6B and Supplementary Figure 5B). ACLY deletion reduced H3K9ac abundance near genes associated with NK cell cytotoxicity (*Prf1*, *Gzmb*, *Gzmc*). Focusing on the genes encoding DAP10 and DAP12, there was a decreased H3K9ac marks within the promoter region of *Tyrobp* (DAP12) gene in KO cells with log_2_FC of –0.58 and FDR of 8.97E-03 (Figure 6C). Interestingly, H3K27ac marks in the same region was increased in ACLY KO (log_2_FC = 1.1, FDR = 1.88E-08) (Figure 6C). In *Hcst* (DAP10) gene, H3K27ac modifications were increased in ACLY KO cells with a log_2_FC of 0.6 and 0.7 (FDR= 5.81E-04 and 1.26E-03 respectively) (Figure 6D), associated with higher transcript and protein expression of DAP10 (Figure 5A). This indicates that ACLY-generated acetyl-CoA contributed to selective histone acetylation of genes encoding adaptor proteins. Moreover, WT cells showed higher H3K9ac marks near the 5’end of NKG2D-S (Supplementary Figure 5C). This suggests that transcription of *Klrk1* isoform may be regulated by histone acetylation.

**Figure 6.**
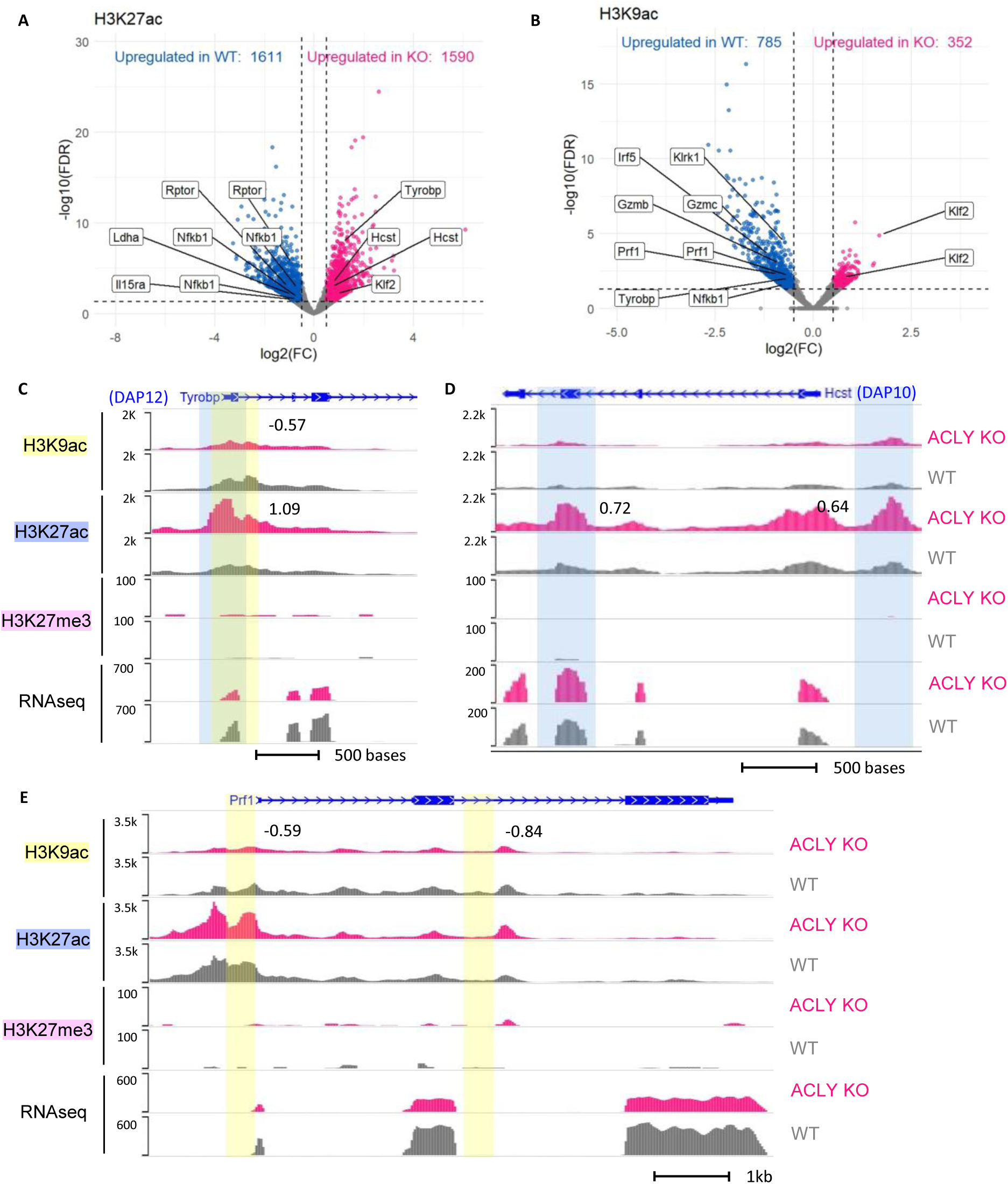
CUT&Tag analysis identifies differential distribution of histone acetylation marks between primed WT and ACLY KO NK cells. Sorted YFP^+^ WT or ACLY KO NK cells were pooled from 3-5 mice for each condition and primed with HD15 for 72 hours followed by CUT&Tag. Results are from n=3-4 biologic replicates for each histone mark and genotype. (**A-B**) Volcano plots showing H3K27ac (A) and H3K9ac (B) signals upregulated in primed YFP^+^ WT or ACLY KO cells. Peaks that were annotated to promoter, exon, or intron are labeled with the assigned gene name. (**C through E**) Genome browser view of H3K9ac, H3K27ac, H3K27me3 CUT&Tag and RNAseq at *Tyrobp* (DAP12) (C), *Hcst* (DAP10) (D) and *Prf1* (E) gene loci. Differential peaks identified in the H3K9ac (highlighted in yellow) or H3K27ac (highlighted in blue) marks are shown with number representing log_2_FC between WT and ACLY KO.

Deletion of *Acly* led to limited amount of acetyl group available for histone acetylation. However, CUT&Tag sequencing data revealed stronger H3K9ac and H3K27ac signal at specific loci in ACLY KO NK cells. We compared the annotated genomic regions of the peaks with higher signal in WT and ACLY KO cells. The genomic distribution showed that in ACLY KO cells, the majority of the H3K9ac and H3K27ac peaks were deposited predominantly in promoter regions (Supplementary Figure 5D). This could suggest that ACLY KO cells were allowing a restricted amount of acetyl group on specific gene promoter sites. In other words, during NK cell priming, acetylation in enhancer regions is mostly reliant on ACLY-generated acetyl-CoA.

Consistent with the role of ACLY in acetylation rather than methylation, fewer H3K27me3 peaks with differential signal were detected (Supplementary Figure 5E). Some of these genes exhibit the expected correlation of activating and repressive histone modification with gene expression. For example, in ACLY KO cells, higher H3K27me3 and lower H3K9ac modifications at *Irf4*, *Irf5* and *Mycn* genes were associated with transcriptional repression, whereas lower H3K27me3 and higher histone acetylation at the *Klf2* and *Tcf7* genes were associated with active transcription.

### Acetate supplementation can rescue functional defects in ACLY KO NK cells

When active, ACLY can support glycolysis through production of NAD^+^, and generation of cytosolic acetyl-CoA (Supplementary Figure 1A). Cytosolic acetyl-CoA is important for reprogramming the epigenome through histone acetylation and can also play a role in protein acetylation and lipid synthesis. An alternative source of cytosolic acetyl-CoA is acetate, which can be processed by the enzyme acyl-CoA synthetase short-chain family member 2 (ACSS2),^47^ and in some cases ACLY and ACSS2 can compensate for each other to maintain cellular functions.^48^ Consequently, acetate supplementation can rescue the cytosolic acetyl-CoA function of ACLY deficiency. For example, CD8^+^ effector T cells switch from glucose to acetate to produce cytosolic acetyl-CoA in the absence of ACLY,^22^ and can rescue impaired function of CD8^+^ T cell in glucose-restricted environment.^49^ Therefore, we tested whether functional defects of primed ACLY KO NK cells are due to the cytosolic acetyl-CoA function of ACLY by supplementing the cells with acetate.

WT and ACLY KO NK cells were primed with HD15 in the presence of acetate (NaOAc, 5mM) which completely rescued proliferation of ACLY KO NK cells (Figure 7A-B) along with upregulated GLUT1 expression (Supplementary Figure 6A). Moreover, acetate supplementation in ACLY KO cells upregulated the level of neutral lipid (Supplementary Figure 6B). NK cells primed in the presence of acetate, washed and then activated with anti-NKG2D or anti-Ly49H had restored IFN-γ production (Figure 7C and Supplementary Figure 6D) and surface expressions of NKG2D and Ly49H (Supplementary Figure 6D-E). Consistent with restored cytokine production, DAP10 and DAP12 protein level also normalized in ACLY KO cells with acetate supplementation (Figure 7D-E, and quantitation in Supplementary Figure 6F-G, 7H-I).

**Figure 7.**
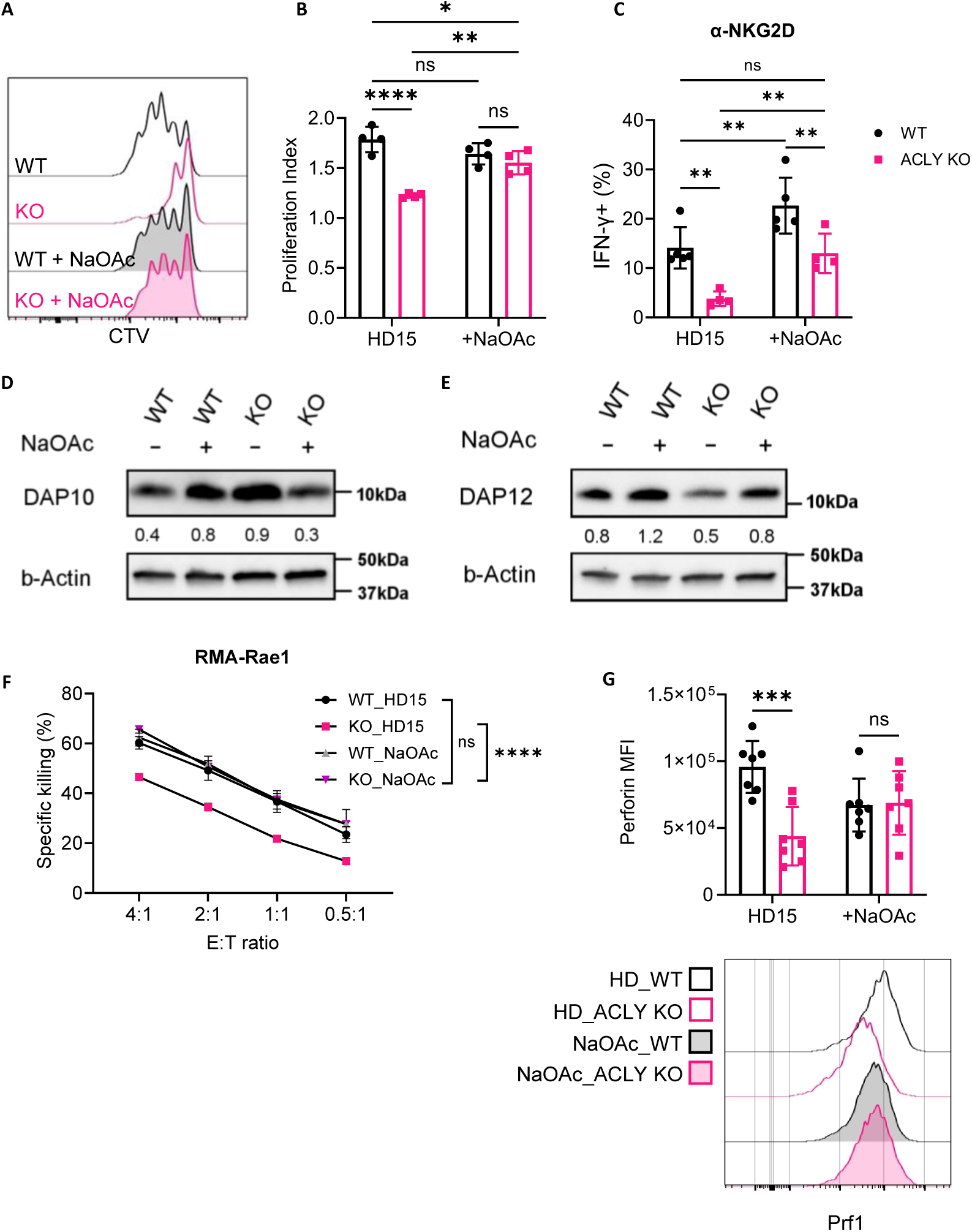
Acetate restores impaired effector function in ACLY KO NK cells. YFP^+^ NK cells were primed with or without 5mM Sodium Acetate (NaOAc) for 72 hours (**A and B**) YFP^+^ NK cells were stained with CTV prior to priming. Representative dilution of CTV of YFP^+^ NK cells (A) and proliferation index (B) shown. Results represent 3 independent experiments with n=4 biological replicates, two-way ANOVA. (**C**) IFN-γ expression in YFP^+^ primed NK cells stimulated with plate-bound NKG2D for 6 hours. Results represent 3 independent experiments with n= 4-5 biological replicates, two-way ANOVA. (**D-E**) Representative protein expression of DAP10 (D) and DAP12 (E) in sorted YFP^+^ NK cells. Protein intensity was quantified by normalizing the intensity of the protein of interest to β-actin. See Figure S7H-I for original image. (**F**) YFP^+^ primed NK cells were incubated with CTV-stained RMA-Rae1 target cells in varying effector-to-target (E:T) ratio for 4 hours. Percent specific killing of targets is shown across E:T ratios. Results represent n=3 sets of pooled mice per group, mean ± SEM, two-way ANOVA. (**G**) MFI and histogram of perforin expression on YFP^+^ NK cells by flow cytometry. Results represent 4 independent experiments with n=7 biological replicates, two-way ANOVA. Data are shown as mean ± SD, *p≤0.05, **p≤0.01, ***p≤0.001, ****p≤0.0001

NK cell cytotoxic capacity was tested after acetate supplementation during priming. Supplementing acetate during priming alone was sufficient to restore the ACLY KO NK cell killing to WT levels against RMA-Rae1 and RMA-m157 (Figure 7F and Supplementary Figure 6H). Consistent with restored cytotoxicity capability, intracellular perforin levels normalized (Figure 7G). Restored killing of targets bearing both NKG2D and Ly49H ligands supports intact signaling through these activating receptors. Together the studies here demonstrate that the ACLY-generated cytosolic acetyl-CoA modulates histone acetylation that is essential for NK cell priming and function

## DISCUSSION

Here, we identified ACLY as a pivotal metabolic enzyme allowing NK cells to modulate their epigenome during priming, thereby enhancing their effector function. While ACLY has been predominantly studied in tumor cells, emerging evidence highlights its critical role in immune cell function and differentiation.^18,19,22–24,50^ Genetic deletion of *Acly* in naïve NK cells does not alter the homeostatic state nor function. However, activation of ACLY KO NK cells with HD15 demonstrates that this enzyme is essential for these innate immune lymphocytes to reprogram to a state of enhanced effector function, including upregulation of glycolysis, proliferation, response to certain activating receptors, and target killing through its role in generating cytosolic acetyl-CoA and histone acetylation.

Interestingly, primed ACLY KO NK cells had a specific impaired response to activating receptors that associate with DAP10 and DAP12, NKG2D and Ly49H, while response to other stimuli including cytokines and an activating receptor that utilizes a different adaptor protein (NK1.1) was completely intact. A prior study had suggested that ACLY was required for NK cell response to cytokines through regulation of OXPHOS,^8^ however this was based on inhibitor data. Here, we show that NK cells genetically deficient in ACLY exhibit normal OXPHOS but impaired glycolysis, and normal response to cytokine stimulation. Our model also highlights that the most commonly used ACLY inhibitor has off target effects, demonstrated by impaired IFN-γ response in ACLY KO cells.

The dynamic process of histone modification is controlled by lysine acetyltransferases (KATs) and lysine deacetylases (KDACs). Differential histone acetylation events are linked to particular KATs.^51^ For example, GCN5/PCAF acetylates H3K9 regions, whereas CBP/p300 acetylates H3K27 regions.^51–53^ This enzymatic activity that changes the epigenome landscape is influenced by the availability of acetyl-CoA, and therefor also by enzymes that facilitate acetyl-CoA production (ACLY, ACSS2, and pyruvate dehydrogenase complex).^42,43,51^ Recruitment of these enzymes to target genes impact the expression of certain genes.^43^ For example, ACLY is recruited to DNA double-strand breaks and ACSS2 is recruited to memory-related genes in hippocampus.^54,55^ Global epigenomic profiling here demonstrates that ACLY deletion alters histone modification levels at specific genes, and as anticipated by the role of ACLY in generate cytosolic acetyl-CoA, primarily affecting acetylation rather than methylation. Investigation of specific targets identified as differentially expressed at the protein level demonstrated that ACLY KO cells had decreased H3K9ac levels near the *Tyrobp* (DAP12) gene, but a stronger H3K27ac marks near the *Hcst* (DAP10) gene. Interestingly, H3K27ac abundance at *Tyrobp* (DAP12) region was elevated in ACLY KO NK cells, potentially compensating for the loss of H3K9ac marks. It is also possible that other histone modifications influence the final gene expression levels.

Differentially expressed DAP10 and DAP12 altered primed NK cell function. NK cell activating receptors, NKG2D and Ly49H, interact with DAP10 and DAP12 through their cytoplasmic tails and interactions with adaptor proteins are critical for cell-surface expression and function of these receptors. In WT NK cells, NKG2D expression showed no difference between primed and naive NK cells, whereas NKG2D was increased on ACLY KO cells, regulated at the transcriptional level. In naïve NK cells, transcript for *Klrk1* (NKG2D) long isoform, which interacts exclusively with DAP10, is expressed. After priming, NK cells generally downregulate transcript for long isoform, while upregulating transcript for short isoform, which interacts with both adaptor proteins. Interestingly, ACLY KO NK cells continued to express high levels of the long isoform while also upregulating transcript for the short isoform. Our CUT&Tag data here demonstrate that WT NK cells showed a higher H3K9ac marks at the region upstream of the *Klrk1* long isoform promoter and between the 5’ end of the short isoform compared to ACLY KO NK cells. As the upstream region of *Klrk1* long isoform promoter has been identified to reduce its activity,^56^ it is possible that transcription changes in NKG2D-L seen in RNA-seq data was likely impacted by epigenetic modifications near the NKG2D-S region. As a result, ACLY KO cells had an overall increase in NKG2D surface protein expression, correlating with abundant DAP10 protein and upregulated expression of NKG2D-L. By contrast, Ly49H primarily associates with DAP12 for surface expression.^5^ Consistent with this, ACLY KO NK cells had lower expression levels of Ly49H associated with less DAP12 protein expression.

ACLY KO NK cells had differences in histone acetylation, but normal intracellular neutral lipids after priming. Of interest, NK cell function in the absence of ACLY was rescued by acetate supplementation, which serves as an alternative source of cytosolic acetyl-CoA that can support histone and protein acetylation as well as lipid synthesis (Supplementary Figure 1A).^48^ When ACLY KO cells were primed with acetate, nearly all phenotypes were completely rescued with the supplementation. The restoration of adaptor proteins, DAP10 and DAP12, and perforin correlated with restored cytokine production and killing of target cells expressing ligands for NKG2D and Ly49H. Acetate also rescued proliferation of ACLY KO NK cells. It is unclear how acetate supplementation promoted NK cells proliferation in our model. Some possibilities include restored GLUT1 expression, as glycolysis is essential for NK cell proliferation.^11,17^ Alternatively or in addition, while ACLY KO NK cells did not have a defect in intracellular lipids after priming, it is possible that the increased lipids with acetate supplementation provided substrate for lipid synthesis including membrane phospholipids.^57^ Additionally, we cannot rule out the contribution of protein acetylation, as cytosolic acetyl-CoA is a key substrate of the process (Supplementary Figure 1A).

In summary, our findings highlight the critical role that the metabolic enzyme ACLY plays in regulating histone acetylation levels in NK cells. Acetyl-CoA generated by ACLY is particularly important during energy-demanding states such as IL-15 priming, and exogenous supplementation with acetate may be a consideration to enhance NK cell effector states. These findings demonstrate a mechanism whereby NK cells are able to rapidly alter their epigenome, in essence hard-wiring enhanced effector responses.

## LIMITATIONS OF THE STUDY

There are several limitations of our study. These experiments demonstrate that ACLY-generated acetyl-CoA regulates the expression of DAP12 in murine NK cells. However, in human NK cells, NKG2D interacts exclusively with DAP10, suggesting that the role of ACLY in human NK cells requires further investigation. Another limitation is that cytosolic acetyl-CoA is important for acetylation of other proteins beyond histones, and we did not comprehensively profile global protein acetylation. Furthermore, the mechanisms by which acetate supplementation rescue some NK phenotypes, in particular proliferation, are not elucidated here, but of significant interest. Finally, while we utilized a genetic model here, we do not know whether *in vivo* supplementation of acetate will alter NK cell function.

## Resource availability

### Lead contact

Requests for further information, resources, and reagents should be directed to and will be fulfilled by the lead contact, Megan A. Cooper.

### Materials availability

No unique materials were generated for this study.

### Data and code availability

RNA and CUT&Tag sequencing data generated during this study are available at Gene Expression Omnibus under accession numbers: GSE286451.

## Acknowledgments

This work was supported by the National Institutes of Health grant R01AI127752 and the CDI Center for Pediatric Immunology at Washington University in St. Louis (to M.A.C.). Editing assistance was provided by InPrint: A Scientific Communication Network at Washington University in St. Louis. We thank Alvin J. Siteman Cancer Center at Washington University School of Medicine in St. Louis for the use of the Siteman Flow Cytometry. The Siteman Cancer Center is supported in part by an NCI Cancer Center Support Grant #P30 CA091842. We would like to acknowledge Eric Tycksen and the Genomic Technology Access Center at the McDonnell Genome Institute at Washington University School of Medicine for help with genomic analysis. We thank Kelsey Trammel, Katherine Owens and Steven Yang for mouse colony management and technical assistance. Some figures were created with BioRender.com.

## Author contributions

Conceptualization, H.S. and M.A.C.; Methodology, H.S., J.M.T., M.A.C.; Investigation, H.S., K.G., J.M.T., J.L., N.S.; Analysis, H.S. A.K., J.T., Y.W.; Writing – original draft, H.S.; Writing – review and editing: M.A.C., and J.E.P.; Visualization, H.S.; Resource, J.E.P.; Supervision, M.A.C., J.E.P., T.A.F.; Funding acquisition, M.A.C.

## Supplemental Information index

Figure S1-S7 and their legends in a PDF

## STAR★Methods

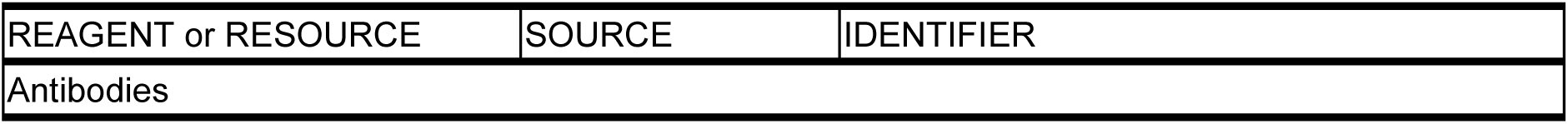
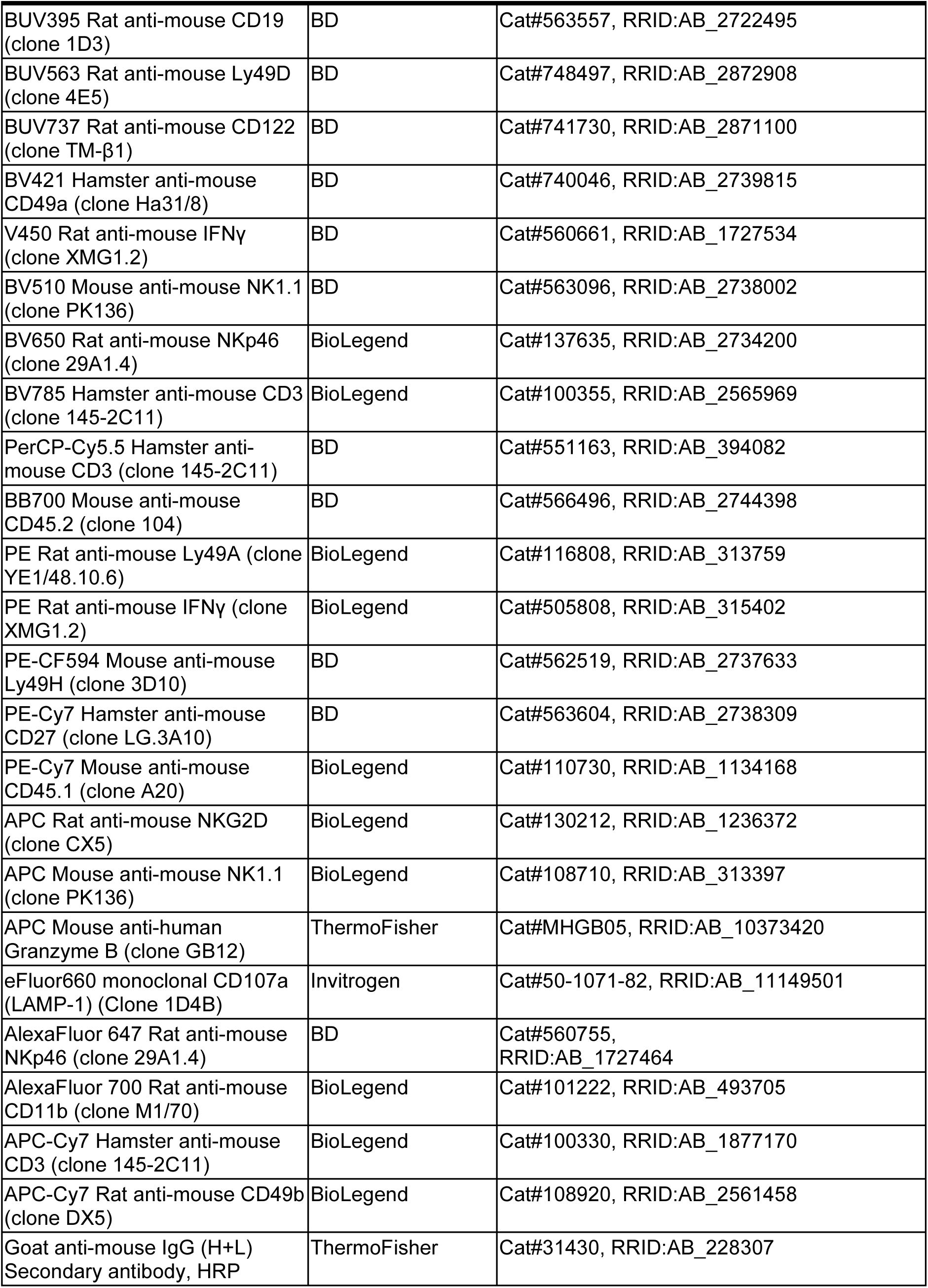
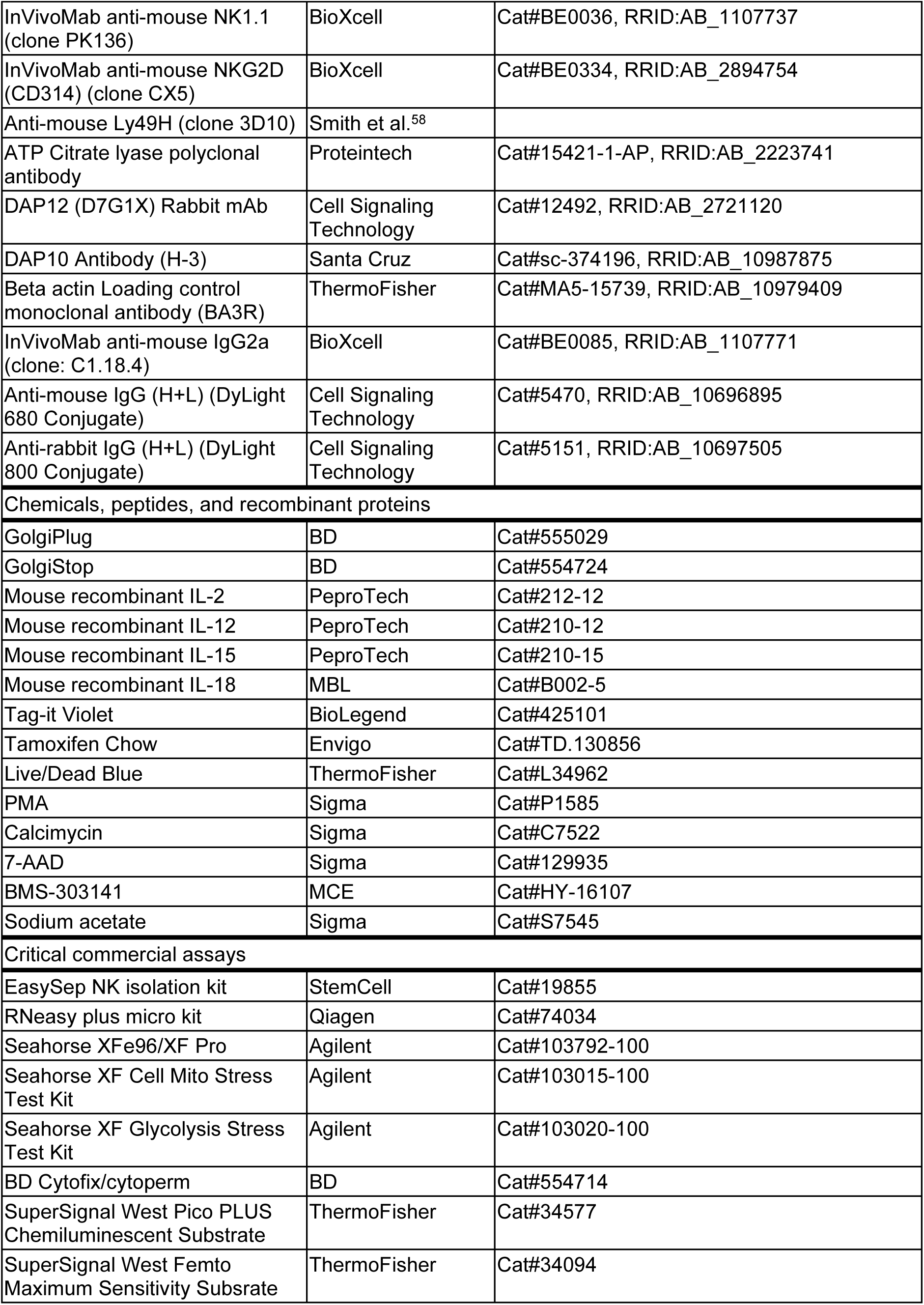
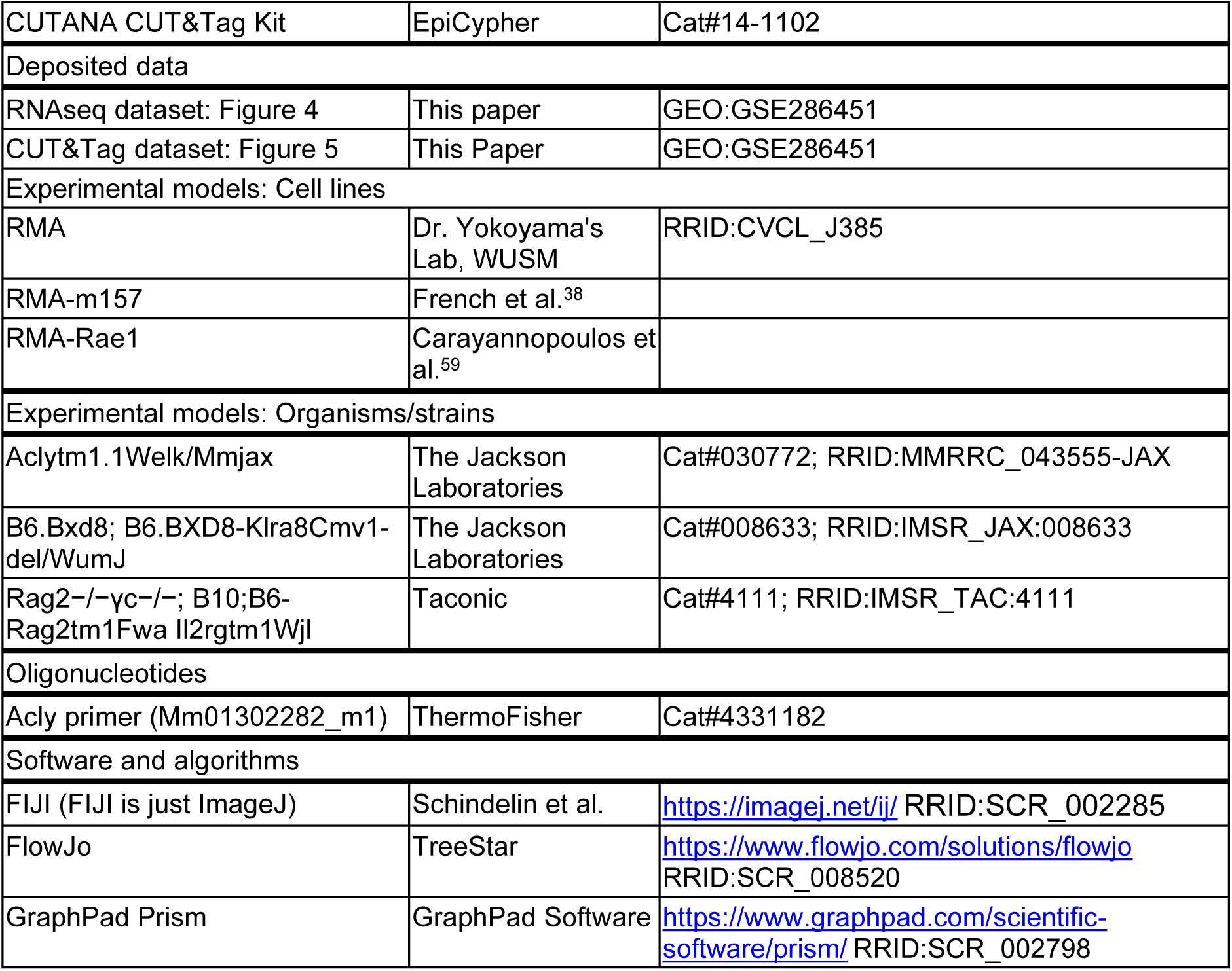
Key resources table.

## Method details

### Mice

An NK cell-specific inducible model of *Acly* deletion (ACLY KO) was generated by crossing mice with tamoxifen-inducible *Ncr1*-restricted Cre expression, *Ncr1*-iCreERT2 mice, that also contain a yellow fluorescent protein (YFP) Cre-reporter (as in Wagner et al.^60^) with mice harboring loxP sites flanking exon 9 of *Acly* (Jackson Laboratory strain #030772).^20^ The resulting mice had tamoxifen-inducible deletion of *Acly* marked by YFP expression, referred to as ‘ACLY KO’ here. CD45.1 or CD45.2 *Ncr1*-iCreERT2 heterozygous mice were used as controls (WT). All mice used in experiments were fed a tamoxifen diet (Inotiv, TD.130856) *ad lib* for 48 hours and then switched to normal diet for 3-5 days prior to analysis. Mice were bred and maintained in specific pathogen-free condition and studies were approved by the Washington University Animal Studies Committee. Experiments were performed on 8-13 weeks old male and female mice.

### NK purification and cell culture

Murine NK cells were enriched from spleens by negative selection (EasySep, StemCell Technologies) and cultured in RPMI 1640 supplemented with 10% FBS, L-glutamine to a final concentration of 4mM, antibiotics (1X Penicillin/Streptomycin), and 50μM 2-Mercaptoethanol (Sigma-Aldrich). For cytokine production, whole splenocytes or enriched NK cells as indicated were stimulated with 10ng/ml murine IL-12 (PeproTech), IL-15 (PeproTech) and 50 ng/ml IL-18 (MBL) for 18 hours, or in plates coated with 20 μg/ml purified antibodies (IgG, NK1.1, NKG2D, or Ly49H) for 6 hours. Brefeldin A (BD) was added to cells after 1 hour of culture and intracellular cytokine production was assayed by flow cytometry using Cytofix/Cytoperm (BD). For priming, enriched NK cells were cultured in 100 ng/ml IL-15 for 72 hours with or without sodium acetate (NaOAc; 5mM). Cells were incubated in 37°C and 5% CO_2_.

### Flow cytometry and cell sorting

A live/dead stain (Invitrogen) was included in most assays to assess cell viability. To prevent nonspecific binding, cells were blocked with anti-FcγRIII (2.4G2) prior to surface staining, which was performed for 15 minutes at 4⁰C. Intracellular staining was conducted using permeabilization buffer following fixation/permeabilization (BD). NK cells were identified by gating on NK1.1 or NKp46. For all flow assays NK cells were gated on YFP^+^ cells. Flow cytometry data were collected on a Cytek Aurora and analyzed using FlowJo software. Cell sorting was performed on a BD FACS Aria II or Cytek Aurora CS sorter. Antibodies: anti-CD107a (1D48), anti-CD11b (M1/70), anti-CD122 (TM-β1), anti-CD19 (1D3), anti-CD27 (LG.3A10), anti-CD3 (145-2C11), anti-CD45.1 (A20), anti-CD45.2 (104), anti-CD49a(Ha31/8), anti-CD49b (DX5), anti-granzyme B (GB12), anti-IFN-γ (XMG1.2), anti-Ly49A (YE1/48.10.6), anti-Ly49D (4E5), anti-Ly49H (3D10), anti-NK1.1(PK136), anti-NKG2D (CX5), anti-NKp46 (29A1.4), anti-perforin (S16009A).

### *In vitro* cytotoxicity assays

Splenocytes from WT and ACLY KO mice were harvested 3 days after the withdrawal of the tamoxifen diet. YFP^+^ NK cells were sorted and primed with or without 5mM NaOAc for 72 hours. Primed (IL-15 cultured) NK cells were mixed with CTV-labeled target cells (RMA, RMA-Rae1, RMA-m157) in the indicated ratios and incubated for 4 hours. Dead cells were identified via 7-AAD staining and specific killing was calculated as: % 7-AAD^+^_effector+target_ – % 7-AAD^+^_target only_.

### Extracellular flux assays

Sorted YFP^+^ cells were primed with 100ng/ml IL-15 for 72 hours with or without 5mM NaOAc. Harvested cells were washed in PBS and resuspended in Seahorse assay media supplemented with L-glutamine (final concentration 2mM) and 1% dialyzed FBS for glycolytic stress test, and additional glucose and sodium pyruvate (final concentration 10mM and 1mM respectively) were supplemented for the mito stress test. 1 x 10^5^ primed cells were plated on poly-L-lysine coated 96-well plate in at least quadruplicate for extracellular flux analysis of oxygen consumption rate (OCR) and extracellular acidification rate (ECAR). Drug condition for Mito Stress Test (Agilent): oligomycin (1μM), FCCP (2μM), Rotenone (0.5μM), and Antimycin A (0.5μM). Drug condition for Glycolytic Stress Test (Agilent): glucose (10mM), oligomycin (1μM), and 2-DG (50mM). Glycolysis rate measurements were calculated as the difference in ECAR measurement before and after glucose injection. Glycolytic capacity measurements were calculated as the difference of ECAR measurement after oligomycin and before glucose injection. Glycolytic reserve was calculated by the difference between glycolytic capacity and glycolysis rate, as described by the manufacturer’s protocol (Agilent, Cat#103020-100).

### Proliferation assays

For IL-15-induced proliferation, NK cells were first labeled with 0.4μM Tag-it Violet (Cell Trace Violet; BioLegend) and cultured in high-dose IL-15 (100ng/ml) for 72 hours. For homeostatic proliferation, YFP^+^ NK cells were sorted from CD45.2 *Ncr1*^iCre-ERT2^*Rosa*^YFP/YFP^*Acly*^fl/fl^ and CD45.1 *Ncr1*^iCre-ERT2^*Rosa*^YFP/YFP^ mice and labeled with 0.4μM Tag-it Violet, mixed at a ratio of 1:1 (ACLY KO to WT) and adoptively transferred into CD45.2 *Rag2*-γc KO mice. NK cells were harvested 3 days after the transfer. Proliferation data were collected on flow cytometry and analyzed using FlowJo software. Proliferation index and percent of proliferated cells are calculated using the proliferation tool in FlowJo software.

### Immunoblotting

Cells were solubilized in lysis buffer (200mM Tris pH6.8, 10% 2-Mercaptoethanol, 8% SDS, 40% Glycerol, 0.4% Bromophenol blue), boiled and sonicated before loading. Primary antibodies used for immunoblotting analysis were monoclonal rabbit anti-DAP12 (Cell Signaling Technology, 12492), mouse anti-DAP10 (Santa Cruz Biotechnology, sc-374196), β-actin monoclonal antibody (ThermoFisher, MA5-15739), anti-ACLY (Proteintech, 15421-1-AP). Cells were loaded on MINI-PROTEAN TGX gels (Bio-Rad, 4569033) and were transferred to PVDF membranes (Millipore, ISEQ20200). Membranes were blocked in 1-5% BSA/TBST for 1 hour at RT. The membranes were incubated with their respective primary antibodies (1:1000) overnight in 4°C. Secondary antibodies (Cell Signaling Technology, 5470, 5151 and ThermoFisher, 31430) for DAP10, DAP12 and β-actin proteins were incubated in 5% skim milk/TBST or TBST for 1 hour at room temperature. For chemiluminescence detection, the membrane was incubated with Pico and Femto substrate (ThermoFisher, 34577, 34094) for 5 min before imaging. Binding was detected by Bio-Rad ChemiDoc Imaging Systems.

### RNA sequencing and analysis

YFP^+^ NK cells were sorted from tamoxifen-treated ACLY KO and WT mice (>95% purity). Cells were primed with high-dose IL-15 for 72 hours, washed and then stimulated with plate-bound anti-NKG2D for an additional 6 hours. Total RNA was isolated from fresh, primed and primed+NKG2D using the RNeasy Micro Plus Kit (QIAGEN). Bulk RNAseq was performed by the Genome Technology Access Center at the McDonnell Genome Institute (GTAC@MGI) at Washington University School of Medicine. Total library preparation was performed with 10 ng of total RNA with a Bioanalyzer RIN score greater than 8.0. cDNA was synthesized with Takara clonetech SMARTer library kit by manufacturer’s protocol and sequenced on an Illumina NovaSeq 6000.

RNA-seq reads were aligned to the Ensembl release 101 primary assembly with an Illumina DRAGEN Bio-IT version 3.9.3-8. All gene counts were imported into EdgeR^61^ and TMM (trimmed mean of M-value) normalization size factors were calculated to adjust for differences in library size. The TMM size factors and the matrix of counts were then imported into the Limma.^62^ Weighted likelihoods based on the observed mean-variance relationship of every gene and sample were calculated for all samples and the count matrix was transformed to moderated log_2_ counts-per-million with Limma’s voomWithQualityWeights.^63^ Differential expression analysis was then performed to identify differences between conditions and the results were filtered for only those genes with Benjamini-Hochberg false-discovery rate adjusted p-values ≤ 1e-5. Bubble plots and GSEA^64^ plots were generated in R with the enrichplot^65^ package. Data has been deposited to Gene Expression Omnibus (GEO) under accession number GSE286451 (https://www.ncbi.nlm.nih.gov/geo/query/acc.cgi?acc=GSE286451).

### CUT&Tag sequencing and analysis

Splenocytes from 3-5 tamoxifen-treated WT or KO mice were pooled together for each replicate. Sorted YFP^+^ NK cells were primed as above, and assessed for H3K27ac, H3K9ac and H3K27me3 abundance using the CUTANA CUT&Tag kit (EpiCypher). For each sample, 100,000 primed cells were used. Libraries were prepared with indexing primers included in EpiCypher kit, cleaned up with SPRIbeads. The resulting libraries were submitted to the GTAC@MGI at Washington University School of Medicine for sequencing on an Illumina NovaSeq X Plus. Data analysis was adapted from Zheng et al.^66^ with the following modifications: The adaptor sequences were removed by Cutadapt before alignment. Trimmed CUT&Tag reads were aligned and mapped to mm10 genome by Bowtie2 software. Peak calling was conducted by MACS2^67^ software. H3K27ac and H3K9ac samples used narrowpeak option with p-value 1e-6 and H3K27me3 samples used broadpeak option with p-value 1e-5. Differential binding analysis was performed by using the Diffbind^68^ R package. BigWig files of same conditions/genotypes were merged using ucsc-bigwigmerge. Data has been deposited in NCBI GEO under accession number GSE286451 (https://www.ncbi.nlm.nih.gov/geo/query/acc.cgi?acc=GSE286451).

### Quantification and statistical analysis

GraphPad Prism 10 was used to analyze all statistics, the detail of which can be found in the figure legends. All statistical data were showed as mean ± SD unless specified. All experiments were independently repeated 2-6 times as indicated. A two-tailed paired or unpaired *t* test was used for comparison of two. For comparisons between genotype and conditions, two-way ANOVA with Sidak’s multiple comparison test. *P < 0.05, **P < 0.01, ***P < 0.001, ****P < 0.0001

## References Cited

1. Pegram, H.J., Andrews, D.M., Smyth, M.J., Darcy, P.K., and Kershaw, M.H. (2011). Activating and inhibitory receptors of natural killer cells. Immunol Cell Biol 89, 216–224. 10.1038/icb.2010.78.

2. Vivier, E., Nunes, J.A., and Vely, F. (2004). Natural killer cell signaling pathways. Science 306, 1517–1519. 10.1126/science.1103478.

3. Gross, O., Grupp, C., Steinberg, C., Zimmermann, S., Strasser, D., Hannesschlager, N., Reindl, W., Jonsson, H., Huo, H., Littman, D.R., et al. (2008). Multiple ITAM-coupled NK-cell receptors engage the Bcl10/Malt1 complex via Carma1 for NF-kappaB and MAPK activation to selectively control cytokine production. Blood 112, 2421–2428. 10.1182/blood-2007-11-123513.

4. Lanier, L.L. (2009). DAP10– and DAP12-associated receptors in innate immunity. Immunol Rev 227, 150–160. 10.1111/j.1600-065X.2008.00720.x.

5. Orr, M.T., Sun, J.C., Hesslein, D.G., Arase, H., Phillips, J.H., Takai, T., and Lanier, L.L. (2009). Ly49H signaling through DAP10 is essential for optimal natural killer cell responses to mouse cytomegalovirus infection. J Exp Med 206, 807–817. 10.1084/jem.20090168.

6. Wensveen, F.M., Jelencic, V., and Polic, B. (2018). NKG2D: A Master Regulator of Immune Cell Responsiveness. Front Immunol 9, 441. 10.3389/fimmu.2018.00441.

7. Colucci, F., Di Santo, J.P., and Leibson, P.J. (2002). Natural killer cell activation in mice and men: different triggers for similar weapons? Nat Immunol 3, 807–813. 10.1038/ni0902-807.

8. Assmann, N., O’Brien, K.L., Donnelly, R.P., Dyck, L., Zaiatz-Bittencourt, V., Loftus, R.M., Heinrich, P., Oefner, P.J., Lynch, L., Gardiner, C.M., et al. (2017). Srebp-controlled glucose metabolism is essential for NK cell functional responses. Nat Immunol 18, 1197–1206. 10.1038/ni.3838.

9. Donnelly, R.P., Loftus, R.M., Keating, S.E., Liou, K.T., Biron, C.A., Gardiner, C.M., and Finlay, D.K. (2014). mTORC1-dependent metabolic reprogramming is a prerequisite for NK cell effector function. J Immunol 193, 4477–4484. 10.4049/jimmunol.1401558.

10. Loftus, R.M., Assmann, N., Kedia-Mehta, N., O’Brien, K.L., Garcia, A., Gillespie, C., Hukelmann, J.L., Oefner, P.J., Lamond, A.I., Gardiner, C.M., et al. (2018). Amino acid-dependent cMyc expression is essential for NK cell metabolic and functional responses in mice. Nat Commun 9, 2341. 10.1038/s41467-018-04719-2.

11. Marcais, A., Cherfils-Vicini, J., Viant, C., Degouve, S., Viel, S., Fenis, A., Rabilloud, J., Mayol, K., Tavares, A., Bienvenu, J., et al. (2014). The metabolic checkpoint kinase mTOR is essential for IL-15 signaling during the development and activation of NK cells. Nat Immunol 15, 749–757. 10.1038/ni.2936.

12. Schafer, J.R., Salzillo, T.C., Chakravarti, N., Kararoudi, M.N., Trikha, P., Foltz, J.A., Wang, R., Li, S., and Lee, D.A. (2019). Education-dependent activation of glycolysis promotes the cytolytic potency of licensed human natural killer cells. J Allergy Clin Immunol 143, 346–358 e346. 10.1016/j.jaci.2018.06.047.

13. Cimpean, M., Keppel, M.P., Gainullina, A., Fan, C., Sohn, H., Schedler, N.C., Swain, A., Kolicheski, A., Shapiro, H., Young, H.A., et al. (2023). IL-15 Priming Alters IFN-gamma Regulation in Murine NK Cells. J Immunol 211, 1481–1493. 10.4049/jimmunol.2300283.

14. Sohn, H., and Cooper, M.A. (2023). Metabolic regulation of NK cell function: implications for immunotherapy. Immunometabolism (Cobham) 5, e00020. 10.1097/IN9.0000000000000020.

15. Keppel, M.P., Saucier, N., Mah, A.Y., Vogel, T.P., and Cooper, M.A. (2015). Activation-specific metabolic requirements for NK Cell IFN-gamma production. J Immunol 194, 1954–1962. 10.4049/jimmunol.1402099.

16. Mah, A.Y., and Cooper, M.A. (2016). Metabolic Regulation of Natural Killer Cell IFN-gamma Production. Crit Rev Immunol 36, 131–147. 10.1615/CritRevImmunol.2016017387.

17. Mah, A.Y., Rashidi, A., Keppel, M.P., Saucier, N., Moore, E.K., Alinger, J.B., Tripathy, S.K., Agarwal, S.K., Jeng, E.K., Wong, H.C., et al. (2017). Glycolytic requirement for NK cell cytotoxicity and cytomegalovirus control. JCI Insight 2. 10.1172/jci.insight.95128.

18. Zaidi, N., Swinnen, J.V., and Smans, K. (2012). ATP-citrate lyase: a key player in cancer metabolism. Cancer Res. 72, 3709–3714. 10.1158/0008-5472.CAN-11-4112.

19. Dominguez, M., Brune, B., and Namgaladze, D. (2021). Exploring the Role of ATP-Citrate Lyase in the Immune System. Front. Immunol. 12, 632526. 10.3389/fimmu.2021.632526.

20. Wellen, K.E., Hatzivassiliou, G., Sachdeva, U.M., Bui, T.V., Cross, J.R., and Thompson, C.B. (2009). ATP-citrate lyase links cellular metabolism to histone acetylation. Science 324, 1076–1080. 10.1126/science.1164097.

21. Peng, M., Yin, N., Chhangawala, S., Xu, K., Leslie, C.S., and Li, M.O. (2016). Aerobic glycolysis promotes T helper 1 cell differentiation through an epigenetic mechanism. Science 354, 481–484. 10.1126/science.aaf6284.

22. Kaymak, I., Watson, M.J., Oswald, B.M., Ma, S., Johnson, B.K., DeCamp, L.M., Mabvakure, B.M., Luda, K.M., Ma, E.H., Lau, K., et al. (2024). ACLY and ACSS2 link nutrient-dependent chromatin accessibility to CD8 T cell effector responses. J Exp Med 221. 10.1084/jem.20231820.

23. Ma, S., Dahabieh, M.S., Mann, T.H., Zhao, S., McDonald, B., Song, W.S., Chung, H.K., Farsakoglu, Y., Garcia-Rivera, L., Hoffmann, F.A., et al. (2024). Nutrient-driven histone code determines exhausted CD8(+) T cell fates. Science, eadj3020. 10.1126/science.adj3020.

24. Osinalde, N., Mitxelena, J., Sanchez-Quiles, V., Akimov, V., Aloria, K., Arizmendi, J.M., Zubiaga, A.M., Blagoev, B., and Kratchmarova, I. (2016). Nuclear Phosphoproteomic Screen Uncovers ACLY as Mediator of IL-2-induced Proliferation of CD4+ T lymphocytes. Mol. Cell. Proteomics 15, 2076–2092. 10.1074/mcp.M115.057158.

25. Gill, K.S., Fernandes, P., O’Donovan, T.R., McKenna, S.L., Doddakula, K.K., Power, D.G., Soden, D.M., and Forde, P.F. (2016). Glycolysis inhibition as a cancer treatment and its role in an anti-tumour immune response. Biochim Biophys Acta 1866, 87–105. 10.1016/j.bbcan.2016.06.005.

26. Hanai, J.I., Doro, N., Seth, P., and Sukhatme, V.P. (2013). ATP citrate lyase knockdown impacts cancer stem cells in vitro. Cell Death Dis 4, e696. 10.1038/cddis.2013.215.

27. Cooper, M.A., Bush, J.E., Fehniger, T.A., VanDeusen, J.B., Waite, R.E., Liu, Y., Aguila, H.L., and Caligiuri, M.A. (2002). In vivo evidence for a dependence on interleukin 15 for survival of natural killer cells. Blood 100, 3633–3638. 10.1182/blood-2001-12-0293 2001-12-0293 [pii].

28. Wagner, J.A., Rosario, M., Romee, R., Berrien-Elliott, M.M., Schneider, S.E., Leong, J.W., Sullivan, R.P., Jewell, B.A., Becker-Hapak, M., Schappe, T., et al. (2017). CD56bright NK cells exhibit potent antitumor responses following IL-15 priming. J Clin Invest 127, 4042–4058. 10.1172/JCI90387.

29. Fehniger, T.A., Cai, S.F., Cao, X., Bredemeyer, A.J., Presti, R.M., French, A.R., and Ley, T.J. (2007). Acquisition of murine NK cell cytotoxicity requires the translation of a pre-existing pool of granzyme B and perforin mRNAs. Immunity 26, 798–811. 10.1016/j.immuni.2007.04.010.

30. Moon, Y.A., Lee, J.J., Park, S.W., Ahn, Y.H., and Kim, K.S. (2000). The roles of sterol regulatory element-binding proteins in the transactivation of the rat ATP citrate-lyase promoter. J Biol Chem 275, 30280–30286. 10.1074/jbc.M001066200.

31. Sato, R., Okamoto, A., Inoue, J., Miyamoto, W., Sakai, Y., Emoto, N., Shimano, H., and Maeda, M. (2000). Transcriptional regulation of the ATP citrate-lyase gene by sterol regulatory element-binding proteins. J Biol Chem 275, 12497–12502. 10.1074/jbc.275.17.12497.

32. Sheppard, S., Srpan, K., Lin, W., Lee, M., Delconte, R.B., Owyong, M., Carmeliet, P., Davis, D.M., Xavier, J.B., Hsu, K.C., and Sun, J.C. (2024). Fatty acid oxidation fuels natural killer cell responses against infection and cancer. Proc Natl Acad Sci U S A 121, e2319254121. 10.1073/pnas.2319254121.

33. Felices, M., Lenvik, A.J., McElmurry, R., Chu, S., Hinderlie, P., Bendzick, L., Geller, M.A., Tolar, J., Blazar, B.R., and Miller, J.S. (2018). Continuous treatment with IL-15 exhausts human NK cells via a metabolic defect. JCI Insight 3. 10.1172/jci.insight.96219.

34. Lunt, S.Y., and Vander Heiden, M.G. (2011). Aerobic glycolysis: meeting the metabolic requirements of cell proliferation. Annu Rev Cell Dev Biol 27, 441–464. 10.1146/annurev-cellbio-092910-154237.

35. Sheppard, S., Santosa, E.K., Lau, C.M., Violante, S., Giovanelli, P., Kim, H., Cross, J.R., Li, M.O., and Sun, J.C. (2021). Lactate dehydrogenase A-dependent aerobic glycolysis promotes natural killer cell anti-viral and anti-tumor function. Cell Rep 35, 109210. 10.1016/j.celrep.2021.109210.

36. Diefenbach, A., Tomasello, E., Lucas, M., Jamieson, A.M., Hsia, J.K., Vivier, E., and Raulet, D.H. (2002). Selective associations with signaling proteins determine stimulatory versus costimulatory activity of NKG2D. Nat Immunol 3, 1142–1149. 10.1038/ni858.

37. Arase, N., Arase, H., Park, S.Y., Ohno, H., Ra, C., and Saito, T. (1997). Association with FcRgamma is essential for activation signal through NKR-P1 (CD161) in natural killer (NK) cells and NK1.1+ T cells. J Exp Med 186, 1957–1963. 10.1084/jem.186.12.1957.

38. French, A.R., Sjolin, H., Kim, S., Koka, R., Yang, L., Young, D.A., Cerboni, C., Tomasello, E., Ma, A., Vivier, E., et al. (2006). DAP12 signaling directly augments proproliferative cytokine stimulation of NK cells during viral infections. J Immunol 177, 4981–4990. 10.4049/jimmunol.177.8.4981.

39. Gilfillan, S., Ho, E.L., Cella, M., Yokoyama, W.M., and Colonna, M. (2002). NKG2D recruits two distinct adapters to trigger NK cell activation and costimulation. Nat Immunol 3, 1150–1155. 10.1038/ni857.

40. Guo, Q., Kang, H., Wang, J., Dong, Y., Peng, R., Zhao, H., Wu, W., Guan, H., and Li, F. (2021). Inhibition of ACLY Leads to Suppression of Osteoclast Differentiation and Function Via Regulation of Histone Acetylation. J Bone Miner Res 36, 2065–2080. 10.1002/jbmr.4399.

41. Lauterbach, M.A., Hanke, J.E., Serefidou, M., Mangan, M.S.J., Kolbe, C.C., Hess, T., Rothe, M., Kaiser, R., Hoss, F., Gehlen, J., et al. (2019). Toll-like Receptor Signaling Rewires Macrophage Metabolism and Promotes Histone Acetylation via ATP-Citrate Lyase. Immunity 51, 997–1011 e1017. 10.1016/j.immuni.2019.11.009.

42. Pietrocola, F., Galluzzi, L., Bravo-San Pedro, J.M., Madeo, F., and Kroemer, G. (2015). Acetyl coenzyme A: a central metabolite and second messenger. Cell Metab 21, 805–821. 10.1016/j.cmet.2015.05.014.

43. Sivanand, S., Viney, I., and Wellen, K.E. (2018). Spatiotemporal Control of Acetyl-CoA Metabolism in Chromatin Regulation. Trends Biochem Sci 43, 61–74. 10.1016/j.tibs.2017.11.004.

44. Kaya-Okur, H.S., Wu, S.J., Codomo, C.A., Pledger, E.S., Bryson, T.D., Henikoff, J.G., Ahmad, K., and Henikoff, S. (2019). CUT&Tag for efficient epigenomic profiling of small samples and single cells. Nat Commun 10, 1930. 10.1038/s41467-019-09982-5.

45. Calo, E., and Wysocka, J. (2013). Modification of enhancer chromatin: what, how, and why? Mol Cell 49, 825–837. 10.1016/j.molcel.2013.01.038.

46. Karmodiya, K., Krebs, A.R., Oulad-Abdelghani, M., Kimura, H., and Tora, L. (2012). H3K9 and H3K14 acetylation co-occur at many gene regulatory elements, while H3K14ac marks a subset of inactive inducible promoters in mouse embryonic stem cells. BMC Genomics 13, 424. 10.1186/1471-2164-13-424.

47. Guertin, D.A., and Wellen, K.E. (2023). Acetyl-CoA metabolism in cancer. Nat Rev Cancer 23, 156–172. 10.1038/s41568-022-00543-5.

48. Zhao, S., Torres, A., Henry, R.A., Trefely, S., Wallace, M., Lee, J.V., Carrer, A., Sengupta, A., Campbell, S.L., Kuo, Y.M., et al. (2016). ATP-Citrate Lyase Controls a Glucose-to-Acetate Metabolic Switch. Cell Rep 17, 1037–1052. 10.1016/j.celrep.2016.09.069.

49. Qiu, J., Villa, M., Sanin, D.E., Buck, M.D., O’Sullivan, D., Ching, R., Matsushita, M., Grzes, K.M., Winkler, F., Chang, C.H., et al. (2019). Acetate Promotes T Cell Effector Function during Glucose Restriction. Cell Rep 27, 2063–2074 e2065. 10.1016/j.celrep.2019.04.022.

50. Noe, J.T., Rendon, B.E., Geller, A.E., Conroy, L.R., Morrissey, S.M., Young, L.E.A., Bruntz, R.C., Kim, E.J., Wise-Mitchell, A., Barbosa de Souza Rizzo, M., et al. (2021). Lactate supports a metabolic-epigenetic link in macrophage polarization. Sci Adv 7, eabi8602. 10.1126/sciadv.abi8602.

51. Shvedunova, M., and Akhtar, A. (2022). Modulation of cellular processes by histone and non-histone protein acetylation. Nat Rev Mol Cell Biol 23, 329–349. 10.1038/s41580-021-00441-y.

52. Jin, Q., Yu, L.R., Wang, L., Zhang, Z., Kasper, L.H., Lee, J.E., Wang, C., Brindle, P.K., Dent, S.Y., and Ge, K. (2011). Distinct roles of GCN5/PCAF-mediated H3K9ac and CBP/p300-mediated H3K18/27ac in nuclear receptor transactivation. EMBO J 30, 249–262. 10.1038/emboj.2010.318.

53. Voss, A.K., and Thomas, T. (2018). Histone Lysine and Genomic Targets of Histone Acetyltransferases in Mammals. Bioessays 40, e1800078. 10.1002/bies.201800078.

54. Sivanand, S., Rhoades, S., Jiang, Q., Lee, J.V., Benci, J., Zhang, J., Yuan, S., Viney, I., Zhao, S., Carrer, A., et al. (2017). Nuclear Acetyl-CoA Production by ACLY Promotes Homologous Recombination. Mol Cell 67, 252–265 e256. 10.1016/j.molcel.2017.06.008.

55. Mews, P., Donahue, G., Drake, A.M., Luczak, V., Abel, T., and Berger, S.L. (2017). Acetyl-CoA synthetase regulates histone acetylation and hippocampal memory. Nature 546, 381–386. 10.1038/nature22405.

56. Lai, C.B., Zhang, Y., Rogers, S.L., and Mager, D.L. (2009). Creation of the two isoforms of rodent NKG2D was driven by a B1 retrotransposon insertion. Nucleic Acids Res 37, 3032–3043. 10.1093/nar/gkp174.

57. Schug, Z.T., Peck, B., Jones, D.T., Zhang, Q., Grosskurth, S., Alam, I.S., Goodwin, L.M., Smethurst, E., Mason, S., Blyth, K., et al. (2015). Acetyl-CoA synthetase 2 promotes acetate utilization and maintains cancer cell growth under metabolic stress. Cancer Cell 27, 57–71. 10.1016/j.ccell.2014.12.002.

58. Smith, H.R., Chuang, H.H., Wang, L.L., Salcedo, M., Heusel, J.W., and Yokoyama, W.M. (2000). Nonstochastic coexpression of activation receptors on murine natural killer cells. J Exp Med 191, 1341–1354. 10.1084/jem.191.8.1341.

59. Carayannopoulos, L.N., Naidenko, O.V., Fremont, D.H., and Yokoyama, W.M. (2002). Cutting edge: murine UL16-binding protein-like transcript 1: a newly described transcript encoding a high-affinity ligand for murine NKG2D. J Immunol 169, 4079–4083. 10.4049/jimmunol.169.8.4079.

60. Wagner, J.A., Wong, P., Schappe, T., Berrien-Elliott, M.M., Cubitt, C., Jaeger, N., Lee, M., Keppel, C.R., Marin, N.D., Foltz, J.A., et al. (2020). Stage-Specific Requirement for Eomes in Mature NK Cell Homeostasis and Cytotoxicity. Cell Rep 31, 107720. 10.1016/j.celrep.2020.107720.

61. Robinson, M.D., McCarthy, D.J., and Smyth, G.K. (2010). edgeR: a Bioconductor package for differential expression analysis of digital gene expression data. Bioinformatics 26, 139–140. 10.1093/bioinformatics/btp616.

62. Ritchie, M.E., Phipson, B., Wu, D., Hu, Y., Law, C.W., Shi, W., and Smyth, G.K. (2015). limma powers differential expression analyses for RNA-sequencing and microarray studies. Nucleic Acids Res 43, e47. 10.1093/nar/gkv007.

63. Liu, R., Holik, A.Z., Su, S., Jansz, N., Chen, K., Leong, H.S., Blewitt, M.E., Asselin-Labat, M.L., Smyth, G.K., and Ritchie, M.E. (2015). Why weight? Modelling sample and observational level variability improves power in RNA-seq analyses. Nucleic Acids Res 43, e97. 10.1093/nar/gkv412.

64. Subramanian, A., Tamayo, P., Mootha, V.K., Mukherjee, S., Ebert, B.L., Gillette, M.A., Paulovich, A., Pomeroy, S.L., Golub, T.R., Lander, E.S., and Mesirov, J.P. (2005). Gene set enrichment analysis: a knowledge-based approach for interpreting genome-wide expression profiles. Proc Natl Acad Sci U S A 102, 15545–15550. 10.1073/pnas.0506580102.

65. Yu, G. (2024). enrichplot: Visualization of Functional Enrichment Result.. 10.18129/B9.bioc.enrichplot.

66. Zheng Y., A.K., Henikoff S. (2020). CUT&Tag Data Processing and Analysis Tutorial.

67. Zhang, Y., Liu, T., Meyer, C.A., Eeckhoute, J., Johnson, D.S., Bernstein, B.E., Nusbaum, C., Myers, R.M., Brown, M., Li, W., and Liu, X.S. (2008). Model-based analysis of ChIP-Seq (MACS). Genome Biol 9, R137. 10.1186/gb-2008-9-9-r137.

68. Stark, R., Brown, G. (2011). DiffBind: differential binding analysis of ChIP-Seq peak data.

