## Supplemental Figure for "ACLY promotes NK cell effector function by regulating glycolysis and histone acetylation"

**Figure S1**

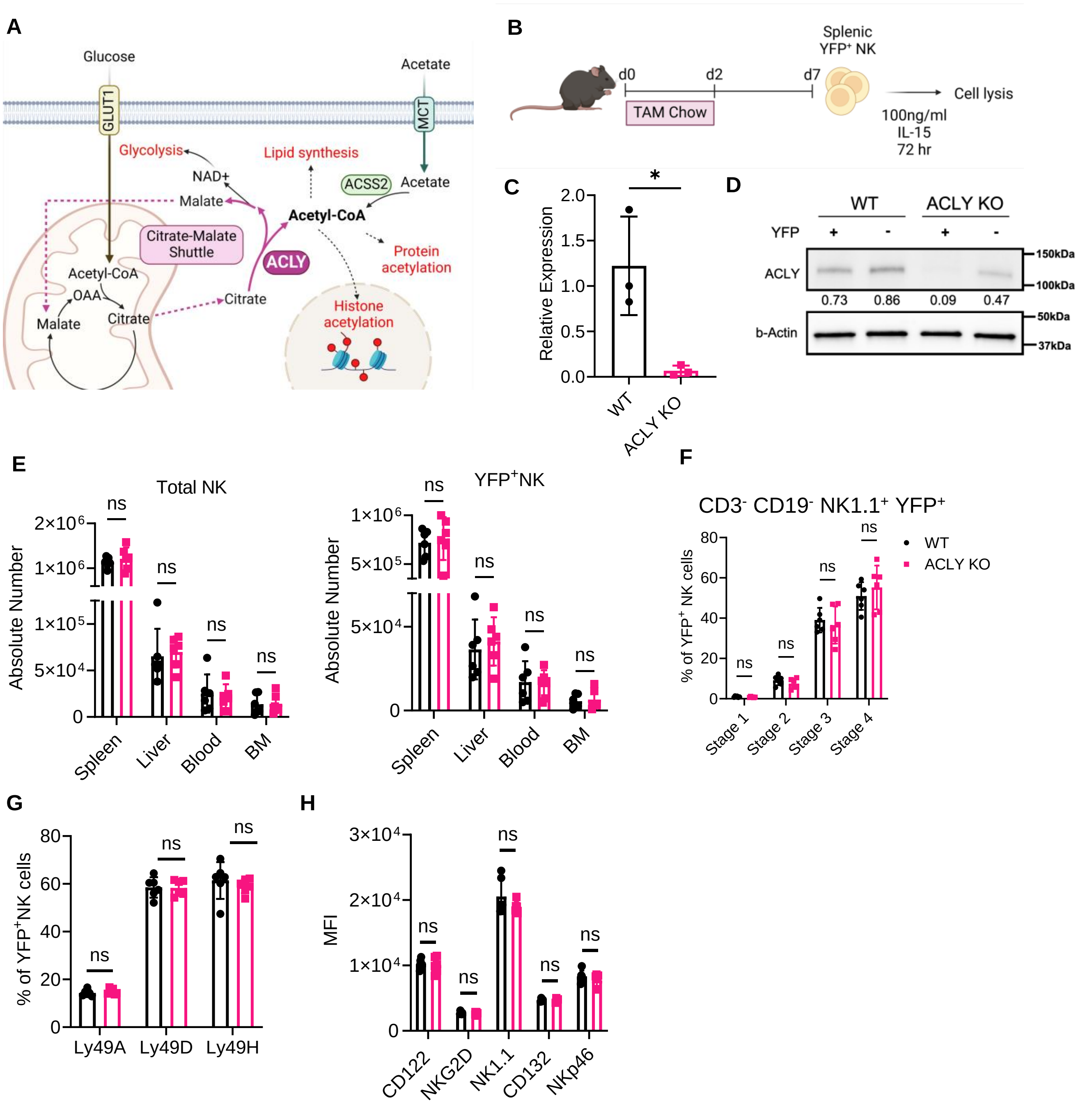


**Supplementary Figure 1. NK cell compartment in ACLY KO mice.** *Rosa*$^{YFP/YFP}$ *Ncr1*$^{iCre\text{-}ERT2}$ (WT) and *Acly*$^{fl/fl}$ *Rosa*$^{YFP/YFP}$ *Ncr1*$^{iCre\text{-}ERT2}$ (ACLY KO) mice were treated with tamoxifen diet for 48 hours. **(A)** ACLY is an enzyme in citrate-malate shuttle that generates cytosolic acetyl-CoA, a substrate for protein and histone acetylation, as well as lipid synthesis. Acetate can serve as another source of cytosolic acetyl-CoA, mediated by enzyme ACSS2. **(B and D)** YFP$^+$ NK cells were sorted 5 days after the last tamoxifen treatment to confirm absence of *Acly* mRNA (C) and protein (D). Unpaired t-test. See Figure S7A-B for original image. **(E)** Total and YFP$^+$ NK cell number in spleen, liver, blood and BM. Two-way ANOVA. **(F)** YFP$^+$ NK cell differentiation was examined by the expression of CD27 and CD11b. (Stage 1: CD27$^-$ CD11b$^-$, Stage 2: CD27$^+$ CD11b$^-$, Stage 3: CD27$^+$ CD11b$^+$, Stage 4: CD27$^-$ CD11b$^+$) NK cells were gated with CD3$^-$CD19$^-$NK1.1$^+$. Two-way ANOVA. **(G and H)** Percentage expression of Ly49 receptor family (G) and Mean Fluorescent Intensity (MFI) of NK receptors (CD122 (IL-15Rβ), NKG2D, NK1.1, CD132 (gc chain) and NKG2D) (H) on YFP$^+$ NK cells. Unpaired t-test. Result represents 2-3 independent experiments with n=4-6 biological replicates. Data are shown as mean ± SD. *p≤0.05, **p≤0.01, ***p≤0.001, ****p≤0.0001

**Figure S2**

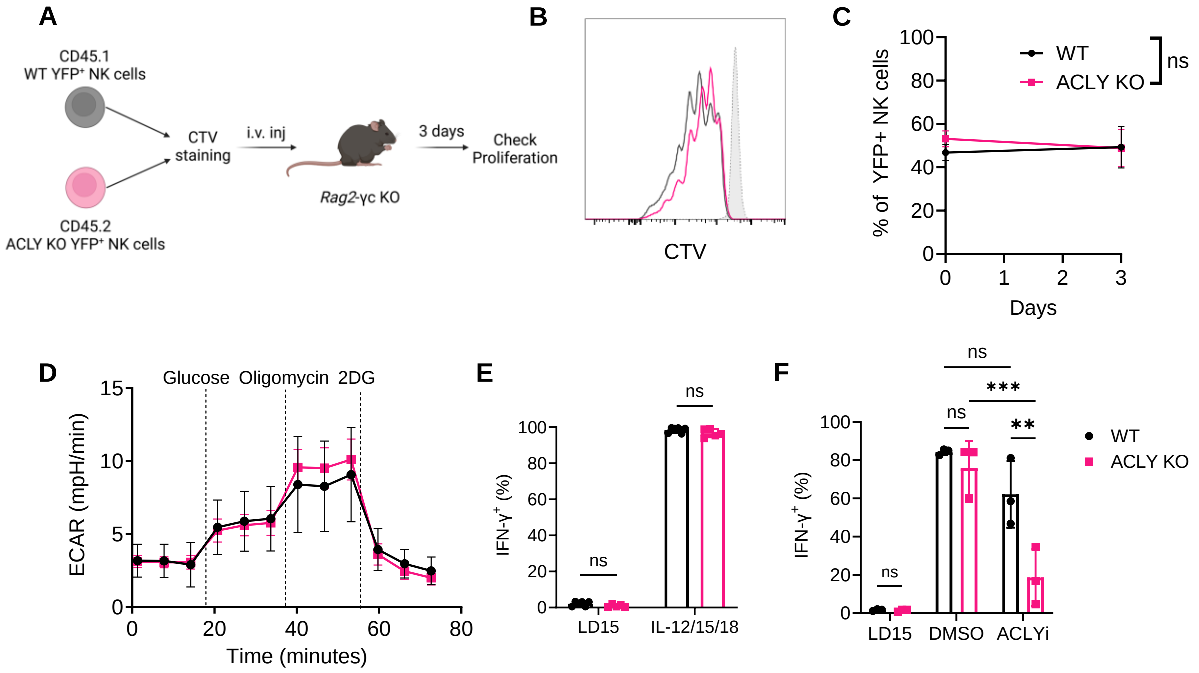


**Supplementary Figure 2. *Acly*-deficient NK cells have intact homeostatic proliferation and cytokine production. (A-C)** (A) YFP$^{+}$ NK cells were sorted from CD45.1 *Rosa*$^{YFP/YFP}$ *Ncr1*$^{iCre\text{-}ERT2}$ (WT) and CD45.2 *Acly*$^{fl/fl}$ *Rosa*$^{YFP/YFP}$ *Ncr1*$^{iCre\text{-}ERT2}$ (ACLY KO) mice. Cells were stained with cell-trace violet (CTV) and adoptively transferred to *Rag2*-gc KO cells in 1:1 ratio. Dilution of CTV (B) and percentages of CD45.1 (WT) and CD45.2 (ACLY KO) within YFP$^{+}$ NK cells (C) were analyzed using flow cytometry. Results represent 2 independent experiments with n=7 biological replicates, paired t-test. **(D)** Extracellular acidification rate (ECAR) of fresh YFP$^{+}$ NK cells from WT and ACLY KO mice. **(E)** IFN-g positive NK cells by intracellular flow cytometry after culture of freshly isolated NK cells with low-dose IL-15 (LD15; 10ng/ml) or IL-12/15/18 (50ng/ml IL-12, 10ng/ml IL-15, 10 ng/ml IL-18) for 18 hours. Results represent 2 independent experiments with n=6 biological replicates, two-way ANOVA. **(F)** Splenocytes were cultured in LD15 for 6 days and stimulated with IL-2 (20 ng/ml) and IL-12 (10 ng/ml) for 18 hours in the presence of DMSO (carrier) or the ACLYi inhibitor BMS-303141 (50mM). Percentage of IFN-g$^{+}$ NK cells was measured by intracellular flow cytometry. Result represents 2 independent experiments with n=3 biological replicates, two-way ANOVA.

**Figure S3**

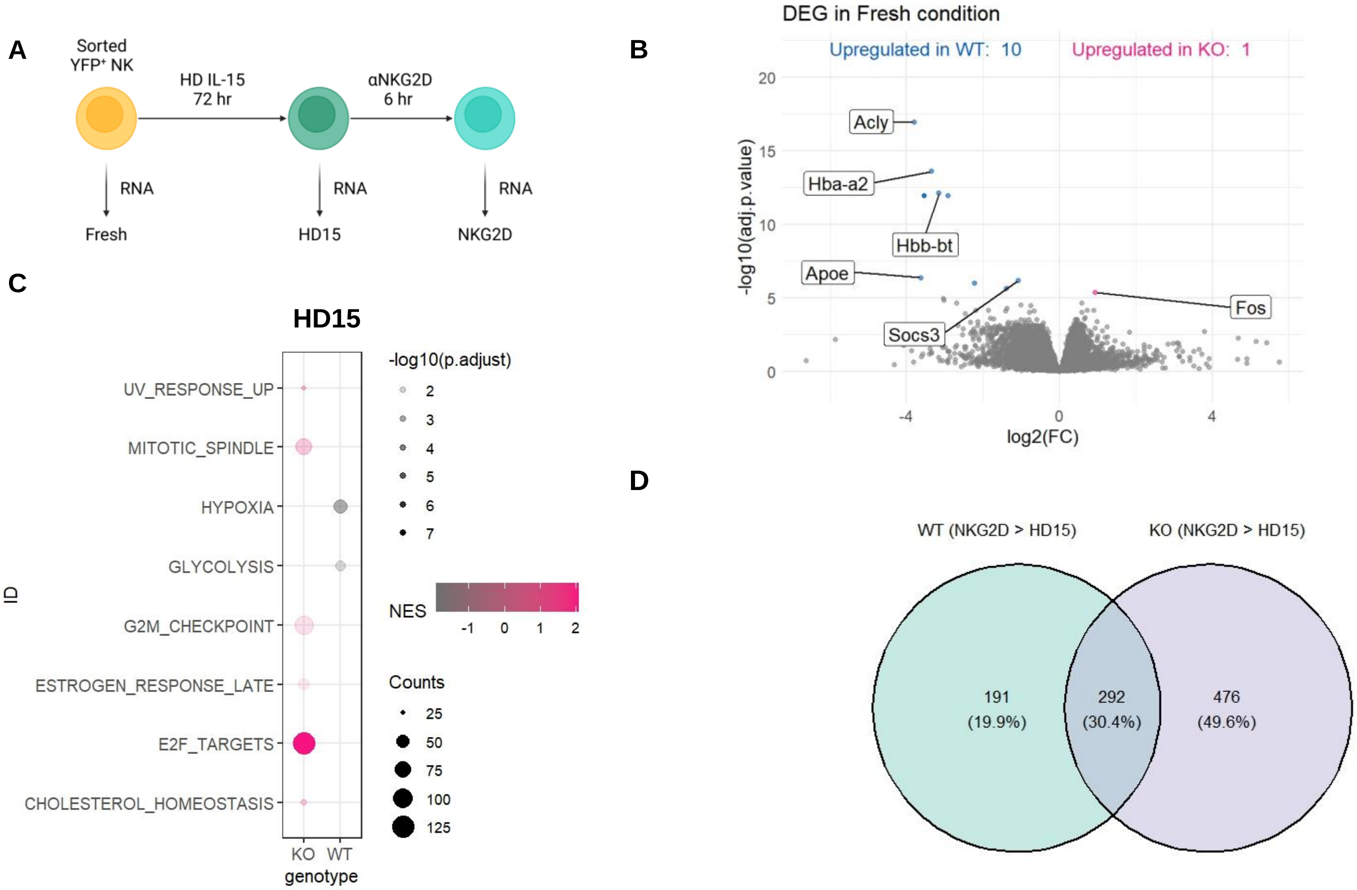


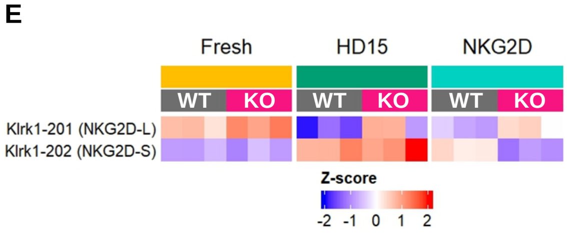


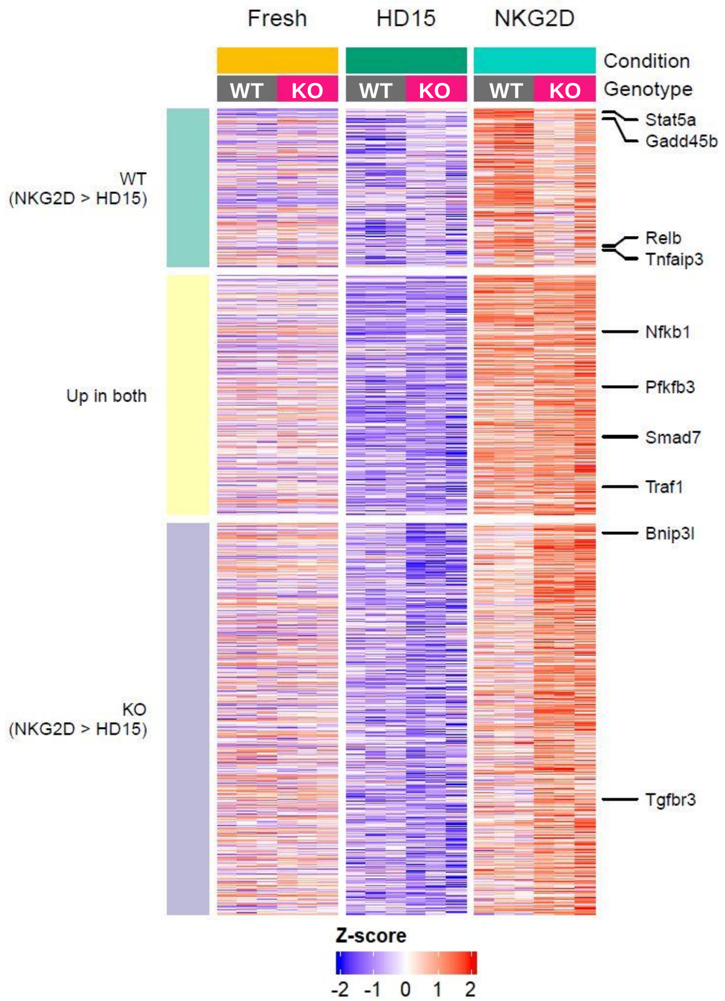


**Supplementary Figure 3. ACLY regulates the transcription of glycolysis-related genes in primed NK cells.** YFP⁺ NK cells from the spleen of 2 mice were pooled for each sample. Results are from one independent experiment with 3 biological replicates per condition and genotype. **(A)** YFP⁺ NK cells are sorted from spleen (Fresh). Fresh NK cells are primed with 100ng/ml IL-15 for 72 hours (HD15). Primed cells were washed and stimulated with plate-bound NKG2D for 6 hours (NKG2D). **(B)** Volcano plots showing differentially expressed genes (DEG) of WT and ACLY KO from the Fresh condition. **(C)** Bubble plot of gene set enrichment analysis (GSEA) of Hallmark pathways upregulated (magenta) and downregulated (gray) in ACLY KO compared to WT in HD15 condition. **(D)** Comparison of genes upregulated in NKG2D over HD15 in WT and KO cells. Genes that had $log_2FC > 0.5$ and adjusted p-value < 1e-5 were called significant. Global heatmap showing all the genes upregulated by NKG2D stimulation. **(E)** Heatmap showing the expression of two NKG2D isoforms (long and short).

**Figure S4**

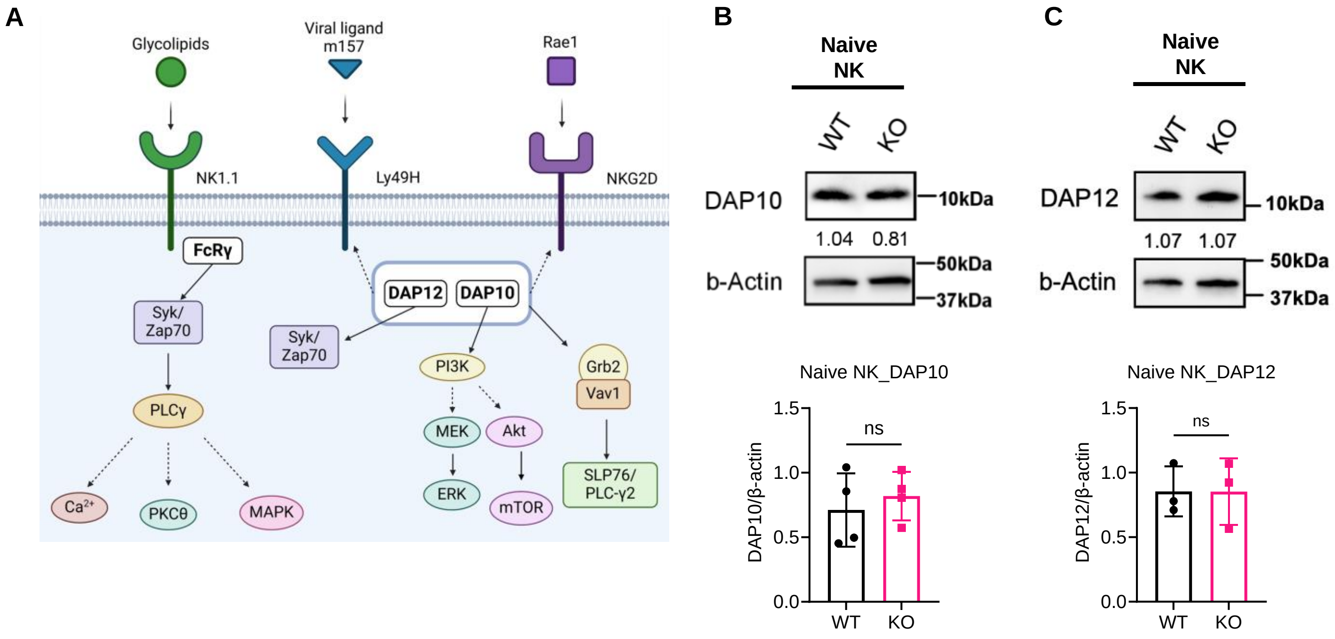


**Supplementary Figure 4. Expression of activating receptor adaptor proteins. (A)** NK1.1 activates Syk/ZAP70 through FcRg. Ly49H and NKG2D are associated with adaptor proteins, DAP10 and DAP12. DAP12 transduce signal to Syk/Zap70, whereas DAP10 activates PI3K and Vav1. **(B and C)** Protein level of DAP10 (B) and DAP12 (C) in naïve NK cells were examined by immunoblotting. Protein intensity was quantified by normalizing the intensity of protein of interest to that of b-actin. Results represent 3-4 independent experiments with n=3-4 biological replicates, paired t-test. See Figure S7F-G for original image.

Figure S5

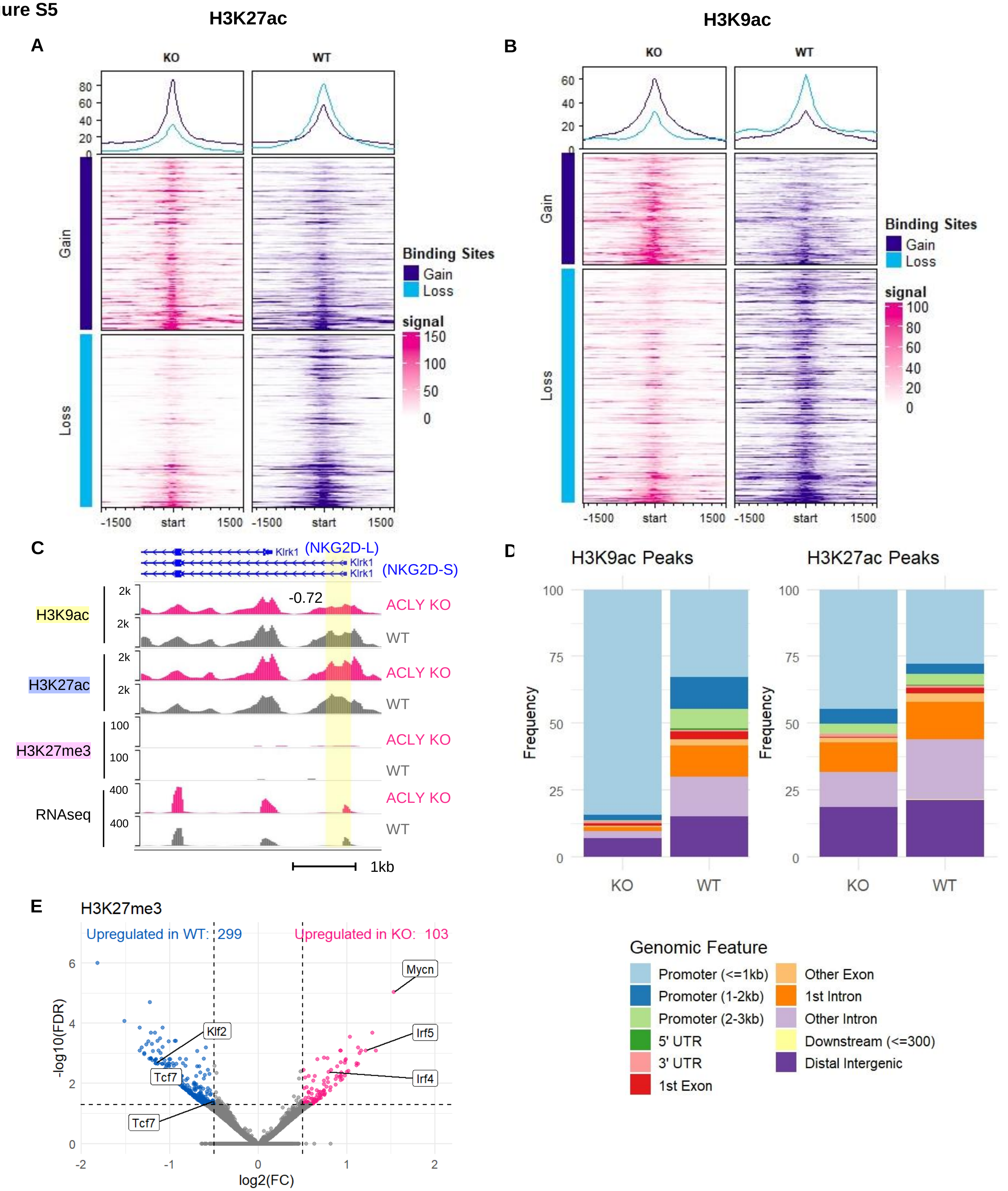


**Supplementary Figure 5. Histone acetylation is reduced in specific regions of ACLY KO NK cells.** YFP⁺ NK cells from the spleen of 3-5 mice were pooled for each sample. NK cells were primed in HD15 for 72 hours. Results are from four independent experiments with 3-4 biological replicates per histone marks and genotype. **(A-B)** Heatmap of H3K27ac (A) and H3K9ac (B) signal across TSS ±1.5kb in genomic regions that are gained or lost signals in KO. **(C)** Genome browser view of H3K9ac, H3K27ac, H3K27me3 and RNAseq at *Klrk1* gene loci. A differential peak identified in the H3K9ac signal (highlighted in yellow) is shown along with its $log_2$FC. **(D)** Genomic distribution of differential H3K9ac and H3K27ac peaks in primed WT and KO NK cells. **(E)** Volcano plots of differentially enriched H3K27me3 marks in primed YFP⁺ NK cells.

**Figure S6**

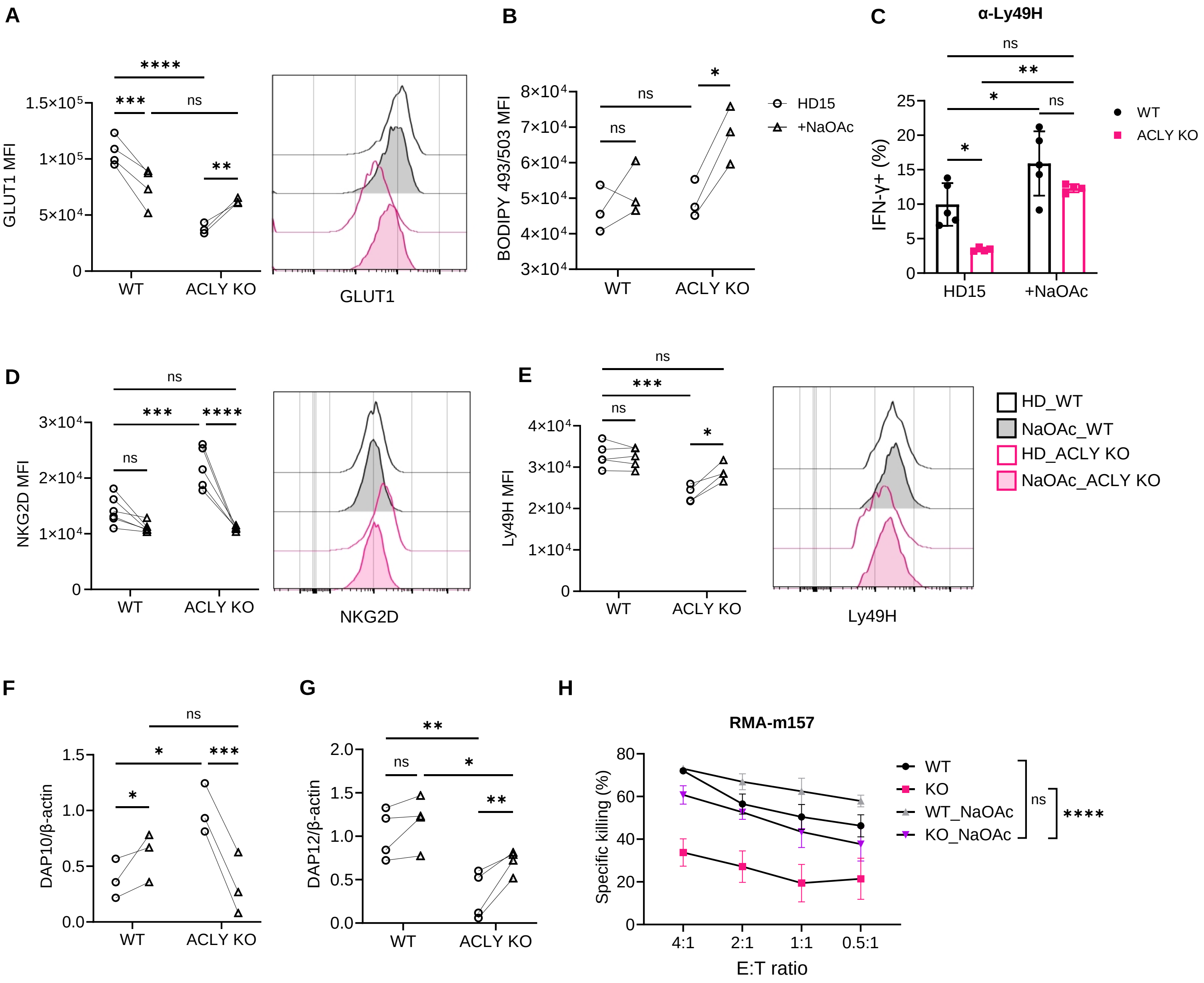


**Supplementary Figure 6. Sodium acetate supplementation partially rescues defects in ACLY KO NK cells.** YFP+ NK cells primed with or without 5mM NaOAc. **(A)** MFI and representative histogram of GLUT1 expression by flow cytometry. Results represent 2 independent experiments with n=4 biological replicates, two-way ANOVA. **(B)** MFI of BODIPY 493/504 level in YFP+ NK cells. Results represent 3 independent experiments with n=3 biological replicates, two-way ANOVA. **(C)** IFN-g expression in primed YFP+ NK cells stimulated with plate-bound Ly49H for 6 hours. Results represent 3 independent experiments with n=5 biological replicates, two-way ANOVA. **(D and E)** MFI and representative histogram of NKG2D on YFP+ (D) and Ly49H on Ly49H+ YFP+ NK cells (E). Results represent 3 independent experiments with n=4-6 biological replicates, two-way ANOVA. **(F and G)** Relative expression levels of DAP10 (F) and DAP12 (G) in primed WT and KO cells. Results represent 3-4 independent experiments, paired t-test. **(H)** YFP+ primed NK cells were incubated with CTV-stained RMA-m157 target cells in indicated effector-to-target (E:T) ratio for 4 hours. Percent specific killing of targets is shown across E:T ratios. Results represent n=3 sets of pooled mice per group, mean ± SEM, two-way ANOVA.

Data are shown as mean ± SD, *p≤0.05, **p≤0.01, ***p≤0.001, ****p≤0.0001

**Figure S7**

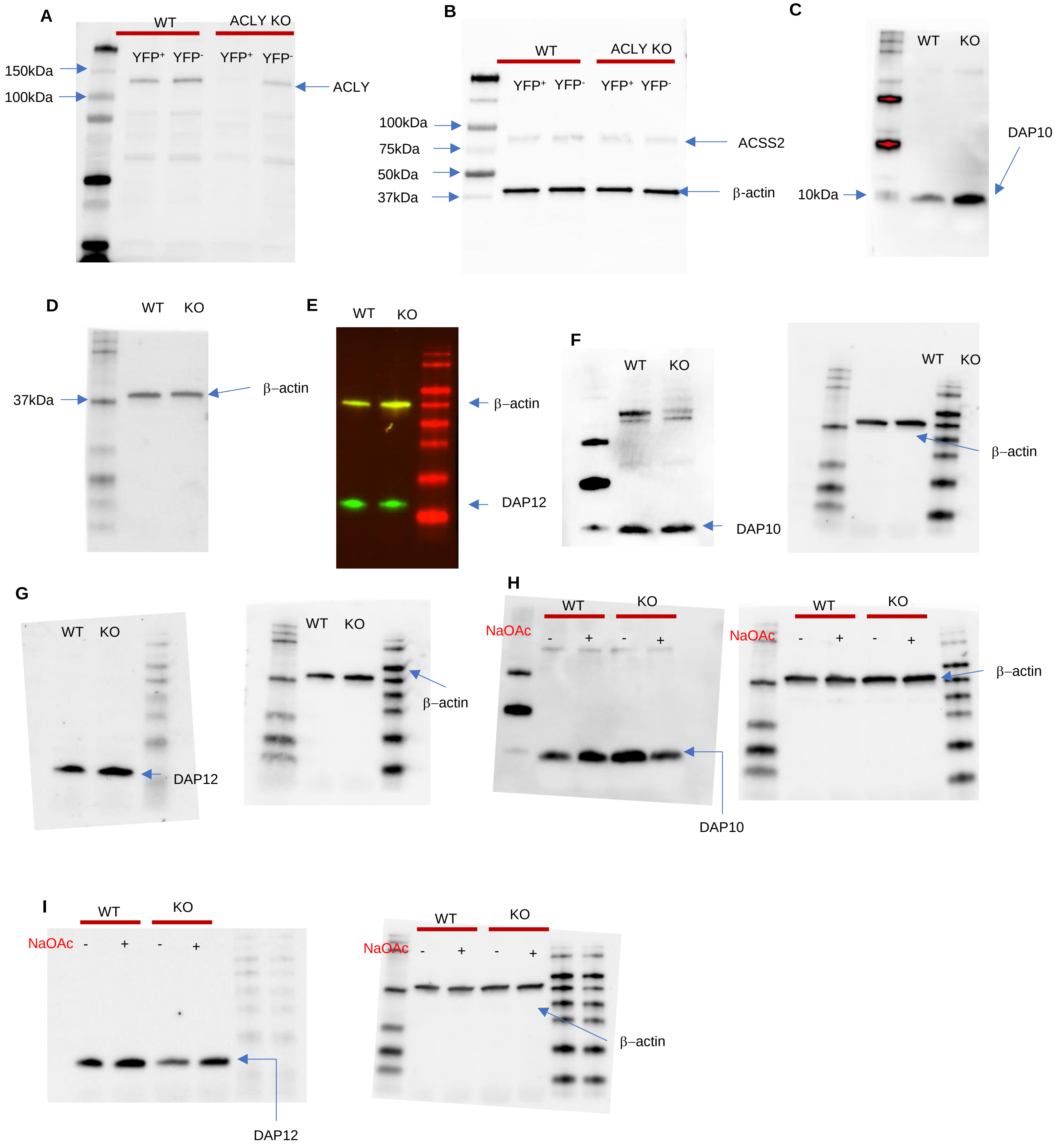
A
WT
ACLY KO
YFP+ YFP- YFP+ YFP-
150kDa
100kDa
ACLY
B
WT
ACLY KO
YFP+ YFP- YFP+ YFP-
100kDa
75kDa
50kDa
37kDa
ACSS2
β-actin
C
WT KO
DAP10
10kDa
D
WT KO
37kDa
β–actin
E
WT KO
β–actin
DAP12
F
WT KO
DAP10
WT KO
β–actin
G
WT KO
DAP12
WT KO
β–actin
H
NaOAc
WT
KO
- + - +
DAP10
NaOAc
WT
KO
- + - +
β–actin
I
NaOAc
WT
KO
- + - +
DAP12
NaOAc
WT
KO
- + - +
β–actin

**Supplementary Figure 7. Original immunoblotting images (A and B)** Confirming ACLY protein deletion in YFP$^+$ cells from ACLY KO mice corresponding to Supplementary Figure 1D. (A) ACLY (120kDa) and (B) b-actin (42kDa) and ACSS2 (78kDa) detected on the same blot. **(C and D)** Original blot corresponding to Figure 5A. (C) DAP10 (10kDa) and (D) b-actin in primed WT and KO cells. **(E)** Original blot corresponding to Figure 5B. DAP12 (12kDa) and b-actin in primed WT and KO cells. **(F and G)** Comparing DAP10 (F), DAP12 (G) and b-actin expression in naive YFP$^+$ NK cells corresponding to Supplementary Figure 4B and 4C respectively. **(H and I)** DAP10 (H), DAP12 (I) and b-actin in primed WT and KO cells without or with 5mM NaOAc, corresponding to Figure 7D and E respectively.
